# Real time, *in vivo* measurement of neuronal and peripheral clocks in *Drosophila melanogaster*

**DOI:** 10.1101/2022.01.12.476067

**Authors:** Peter S. Johnstone, Maite Ogueta, Inan Top, Sheyum Syed, Ralf Stanewsky, Deniz Top

## Abstract

Circadian clocks are highly conserved transcriptional regulators that control 24-hour oscillations in gene expression, physiological function, and behavior. Circadian clocks exist in almost every tissue and are thought to control tissue-specific gene expression and function, synchronized by the brain clock. Many disease states are associated with loss of circadian regulation. How and when circadian clocks fail during pathogenesis remains largely unknown because it is currently difficult to monitor tissue-specific clock function in intact organisms. Here, we developed a method to directly measure the transcriptional oscillation of distinct neuronal and peripheral clocks in live, intact *Drosophila*, which we term Locally Activatable BioLuminescence or LABL. Using this method, we observed that specific neuronal and peripheral clocks exhibit distinct transcription properties. Loss of the receptor for PDF, a circadian neurotransmitter critical for the function of the brain clock, disrupts circadian locomotor activity but not all tissue-specific circadian clocks; we found that, while peripheral clocks in non-neuronal tissues were less stable after the loss of PDF signaling, they continued to oscillate. This result suggests that the presumed dominance of the brain clock in regulating peripheral clocks needs to be re-examined. This result further demonstrates that LABL allows rapid, affordable, and direct real-time monitoring of clocks *in vivo*.

## INTRODUCTION

“Circadian rhythms” collectively refer to 24-hour oscillations in an animal’s behavior and physiological responses. These rhythms are regulated by the circadian clock, transcription/translation negative feedback loops that control 24-hour oscillations in expression of hundreds of genes in every tissue. Circadian clocks are highly evolutionarily conserved timing machines from flies to humans. In both organisms, specialized neurons that express circadian clock components are considered the "central clock"; circadian clock components in non-neuronal tissue (hereafter, "peripheral clocks") are widely assumed to respond to the central clock (Brown et al., 2019; Franco et al., 2018; Ito and Tomioka, 2016; Patke et al., 2020; Pilorz et al., 2018), likely through secreted factors (Handler and Konopka, 1979).

In humans, disruption of the circadian clock is associated with a wide range of pathologies, including neurological, cardiovascular, and metabolic disorders, as well as cancer and aging (Acosta-Rodríguez et al., 2021; Bae et al., 2019; Hood and Amir, 2017a, 2017b; Leng et al., 2019; Logan and McClung, 2019; Rana et al., 2020; Shimizu et al., 2016; Sulli et al., 2019; Thosar et al., 2018; Tsuchiya et al., 2020; Zhang et al., 2021). Such a broad variety of pathologies associated with compromising circadian rhythms suggests a need for cheap and effective ways to measure tissue-specific circadian clocks directly. In animals, locomotion is the simplest and most rapid way to measure circadian clock output, but this output embodies the sum of activity of many clocks and does not necessarily represent all clocks equally. Moreover, while ablation of neuronal clocks in flies, mice, and humans leads to loss of sleep/activity rhythms and is thought to cause loss of circadian clock function in all tissues, the hierarchy of dysfunction of tissue-specific clocks during specific disease pathogenesis remains unclear. Currently, individual peripheral clocks can be measured by removing the organ and extracting RNA to assess transcriptional oscillations (Erion et al., 2016; Gill et al., 2015; Selcho et al., 2017). Such terminal qRT-PCR outputs from explanted organs measure clock function only for that time point in the lifespan of the organism and can be time-, cost-, and labor-intensive. Given that circadian clocks appear to be linked to a wide range of physiologies, including metabolism, as well as various behavioral disorders, there is a need to monitor distinct cell- and tissue-specific circadian clocks directly, *in vivo*, and in real time.

We developed a genetically encoded reporter to monitor distinct clocks in *Drosophila* that we call <u>L</u>ocally <u>A</u>ctivatable <u>B</u>io<u>L</u>uminescence (LABL), offering both high spatial and temporal resolution of clock oscillations *in vivo*. Our data reveals that tissue-specific clocks have similar but distinct properties of oscillation. To determine if tissue-specific clocks are differentially affected by whole-body mutations, we tested flies lacking a functional PDF receptor (*han*^5304^) (Hyun et al., 2005). PDF is a neuropeptide that elicits a cAMP response from most circadian neurons in the brain and is required to maintain rhythmic behaviour in constant dark conditions (Helfrich-Förster, 1995; Helfrich-Förster et al., 2000; Park et al., 2000; Renn et al., 1999; Shafer et al., 2008). While both *han*^5304^ and *tim*^01^ (lacking a functional clock) flies become behaviourally arrhythmic in constant dark conditions, quantification of *han*^5304^ fly neuronal clocks reveals infradian oscillations of ∼60 hours, suggesting that loss of different circadian components can cause loss of circadian locomotor activity in different ways. Interestingly, when peripheral clocks of a *han*^5304^ mutant fly are investigated, they continue to oscillate, but with decreased stability.

Here we demonstrate that LABL reporter flies can be used to measure distinct circadian clocks in different neuronal subpopulations and peripheral tissues in real time and *in vivo*. The differential changes to distinct clocks caused by the *han*^5304^ mutation underscores the assertion that tissue-specific circadian clocks are differentially regulated. We believe that this technology will be critical in the interrogation of distinct circadian clocks in future studies, particularly in monitoring peripheral clock function during disease progression.

## RESULTS

### Construction of LABL

To monitor distinct clock oscillations in real time, *in vivo*, we designed a genetically encoded reporter that we call <u>L</u>ocally <u>A</u>ctivatable <u>B</u>io<u>L</u>uminescence (LABL). LABL is constructed into an attB cloning vector for Drosophila embryo injection and PhiC31-mediated genome integration (Bischof et al., 2007) (Supplementary Figure 1). LABL comprises a *per* promoter fused to mCherry flanked by FRT sequences, which is subsequently fused to Luc2 (pGL4.10) (Figure 1A). Three stop codons placed 3’ of mCherry are designed to block Luciferase expression. The employed *per* promoter (∼6.7 kb) responds to clock regulation through the CLK/CYC transcription activator complex (Bargiello et al., 1984). Tissue-specific expression of Flipase (FLP) triggers recombination at the FRT sites, excising mCherry out of the genome and leaving Luciferase under *per* promoter control. To test the functionality of LABL *in vitro*, we monitored FLP-driven LABL activity in cultured S2 cells (Figures 1B, 1C). Since S2 cells do not express CLK, which is required to activate the *per* promoter, we co-transfected LABL plasmid with *clk*- CFP, with and without *flp*. These cells were subsequently imaged for CFP and mCherry expression and monitored for luminescence. Cells expressing *flp* exhibited increased luminescence and a loss of mCherry fluorescence. Cells lacking *flp* exhibited no luminescence but increased mCherry fluorescence. Thus, these data indicate that the LABL reporter is functional as designed and can be activated by FLP expression.

**Figure 1.**
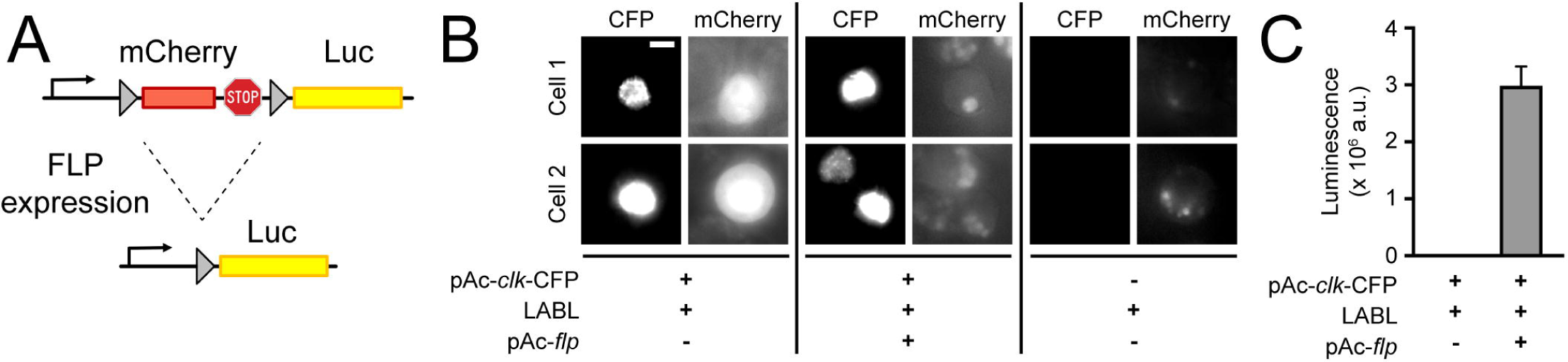
Design and activity of LABL reporter. (**A**) The Locally Activatable Bioluminescence (LABL) reporter construct. In architecture, the *per* promoter is fused to *mCherry* followed by *luciferase* (Luc). The *mCherry* gene, including its three stop codons, is flanked by FRT recombination sequences. Expression of Flipase (FLP) excises out *mCherry* (dashed lines), leaving luciferase under *period* promoter regulation. (**B**) Fluorescence image of LABL-expressing S2 cells. Expression of LABL reporter with *clk* (pAc-*clk*-CFP), *flp* (pAc-*flp*) or both reveal CLK-dependent expression of mCherry, in the absence of FLP. Scale bar represent 5 μm. (**C**) LABL can be activated in cultured S2 cells. Lysed S2 cells emit measurable luminescence is luciferase assay when expressing LABL reporter in a FLP-dependent manner.

### Luminescence oscillations of transcription activity reflect behavioural rhythms

Having established its functionality in cultured cells, we proceeded to assess LABL in adult flies. The LABL reporter plasmid was used to generate reporter flies which were then monitored in a luminometer using arenas designed to hold 15 flies on top of fly food with luciferin (Figure 2A). LABL flies carrying the *tim*-UAS-Gal4 (TUG) pan-circadian tissue driver (Blau and Young, 1999) were crossed to flies carrying UAS-*flp*, and the progeny monitored for luminescence activity in constant darkness (Figure 2B). The raw luminescence data exhibited an oscillating rhythm and a gradual decay, as expected (Brandes et al., 1996; Stanewsky et al., 1997).

**Figure 2.**
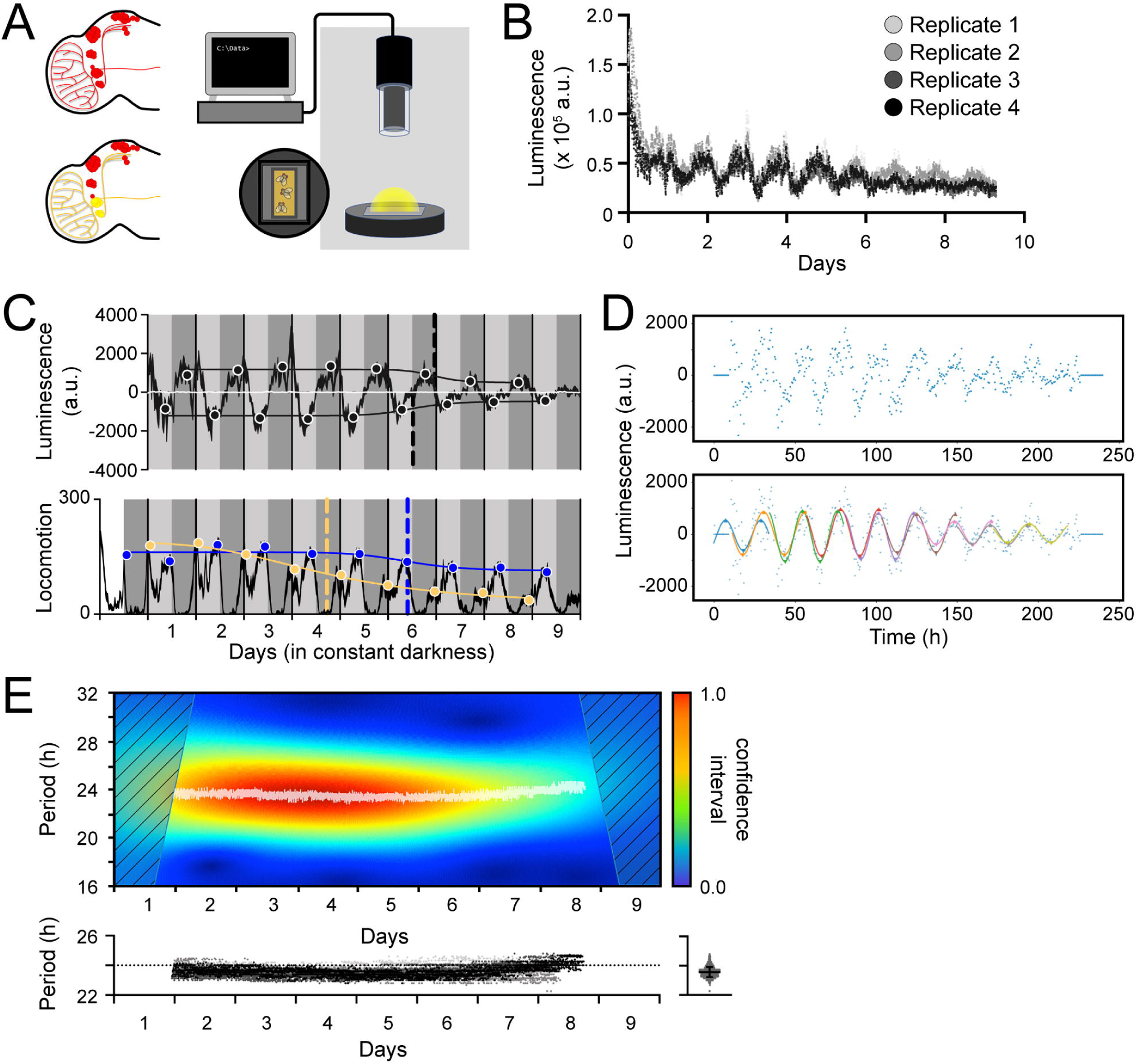
Measurement, quantification and analysis of luminescence from LABL flies. (**A**) LABL activation strategy in *Drosophila* brain. Fly brain schematic illustrates how LABL can be activated using tissue-specific Gal4 drivers to express UAS-FLP2 to excise *mCherry* out of the genome in some neurons to permit Luciferase expression (yellow circles) under the regulation of the *period* promoter, leaving other neurons untouched (red circles). Fifteen LABL flies are placed in custom-made plates containing luciferin mixed with standard fly food. Plates are assembled into a luminometer and luminescence from each cohort is recorded for analysis. (**B**) Raw luminescence measurements of live flies. Photons were detected from four replicates of flies expressing LABL reporter, UAS-FLP2 and *tim*-UAS-Gal4 over 9 days at 4-minute time resolution. (**C**) Luminescence signal and locomotor activity of wild type flies compared. Luminescence signal from flies described in panel B is normalized to exponential decay of signal, the values are averaged into 30-minute bins, and the mean of the four experiments presented, +/- SEM (thickness of curve). Light grey and dark grey backgrounds represent subjective day and subjective night, respectively. Vertical solid black lines divide days. The peaks and troughs of oscillation are represented by a dot are the mean of four experiments, +/- SEM. Line connecting the dots is a best-fit S-curve. The vertical dashed lines define the point of inflection of each S-curve, and is used to define the “amplitude stability constant”. Lower graph represents locomotor activity. Locomotion activity is measured in beam breaks (counts) per 30-minute bouts. Curve represents rhythmic locomotor activity of 25 flies, +/- SEM (thickness of curve). White background indicates lights on. Vertical solid lines divide days. Light grey and dark grey backgrounds represent subjective day and subjective night, respectively. Blue and yellow dots represent peaks of “morning” and “evening anticipatory” locomotion activity, respectively. The vertical dashed lines define the point of inflection of each S-curve, fitted to peaks of activity and is used to define the decay of amplitude of peaks of behavior. (**D**) Calculation of oscillation peaks and troughs. A representative single replicate from experiment in panels B and C is plotted. Dots represent averaged luminescence signal in 30-minute bins. A sinusoidal curve spanning 2 days is fitted to the data in 1-day increments (distinct colored curves). The peaks and troughs of each curve is calculated (triangles), averaged for both x- and y-values and recorded. This process is repeated for all four replicates and the resulting average is reported as shown in panel C. (**E**) Changes in oscillation period over time determined by Morlet wavelet fitting. Wavelets of different periods were fitted to luminescence signal from a single representative replicate from experiment in panel B and C, and assigned a confidence interval, across time (upper graph). Periods with highest confidence intervals at a time point (i.e. across the x-axis) were plotted as white dots. Confidence intervals of 25% or less were omitted. These values were replotted along with the other replicates (below), with varying shades of grey representing each of the four experiments. The dotted horizontal line denotes 24 hours as a point of reference. Right panel: All data points without the time dimension are plotted. Bar represents the mean, +/- SD.

We next compared pan-circadian luminescence oscillations with behavioural rhythms. The recorded luminescence activity was normalized to the gradual decay in signal and plotted as an average of four experiments (Figure 2C, top panel). Control flies lacking a driver exhibited no discernable oscillation (white line). To characterize the luminescence oscillations, we first quantified the decay in amplitude of signal. Data points were binned into 30-minute time intervals and a 48-hour sinusoidal curve was fitted to the data at 24-hour intervals (Figure 2D). The coordinates of the local minima and maxima were recorded, averaged, and plotted over the decay-normalized luminescence signal to reveal the amplitude of oscillation across time (black circles) (Figure 2C). An S-curve fitted to the changing local minima and maxima (black line) revealed points of inflection coinciding to Day 6 of constant darkness (vertical dashed lines). We used this point of inflection as a measure of clock stability (“amplitude stability constant”) since decay of oscillations into arrhythmic transcription may exceed the timeline of this assay.

Genotypically identical flies were measured for locomotor activity and their behavioral rhythms plotted (Figure 2C, bottom panel). We found that the peaks of morning anticipation (yellow circles) decayed rapidly, allowing the evening anticipation peaks (blue circles) to dominate behavioral oscillations in constant dark conditions. Focusing on the change in evening anticipation peaks, we found that a fitted S-curve revealed a point of inflection that falls to ∼ day 6, coincident to the amplitude stability constant observed in luminescence oscillations. We conclude that the decay of oscillation (amplitude) of the molecular clock and behavioural rhythms are consistent with each other.

We next characterized the change of period of luminescence oscillation across time. A Morlet wavelet was fitted onto the measured luminescence oscillations (Figure 2E). Period values with highest confidence intervals revealed a steady ∼23.5 h period of oscillation across time; specifically, oscillations occurred with an average period of 23.57 h over nine days in constant darkness. The decay in oscillation amplitude mirrored the decay in amplitude in behavioral rhythms, with both peaks stabilizing at a lower level on Day 6-7 of constant darkness (Figure 2C). The oscillation period also mirrored the behavioural period, with luminescence oscillation and behavioural period statistically the same, at ∼ 23.5 h. Additionally, luminescence signals peaked at subjective night, when *per* promoter is expected to be active (PER protein peaks at ZT22) and locomotor behavior is expected to be low, oscillating in an approximate reverse phase. Thus, (1) luminescence could be detected in flies in which LABL was activated using UAS-*flp* and a Gal4 driver, and (2) activation of LABL in circadian tissues using the pan-circadian driver TUG revealed parallels between transcription oscillations and behavioral rhythms of the fly.

### LABL signal is comparable to ubiquitously expressed luciferase reporters and is clock dependent

To determine the effectiveness of LABL, we compared TUG-activated LABL oscillations to other, ubiquitously expressed luciferase reporters. To this end, we compared our data to data collected from *plo* (*per* promoter fused to luciferase) (Brandes et al., 1996) and PER-BG::Luc (*per* promoter and ∼2/3rds of the *per* gene fused to luciferase) (Stanewsky et al., 1997) flies, two other luciferase-based reporter systems (Figure 3A). Since TUG drives Gal4 expression in all clock cells, we expected TUG-activated LABL luminescence signal to be comparable to *plo* luminescence signal. Indeed, the oscillation of luminescence signal was comparable in amplitude, period, and phase (Figure 3A). The PER-BG::Luc construct, on the other hand, retains a significant portion of the PER protein, and since there is a phase delay between *per* transcription and PER translation (So and Rosbash, 1997; Stanewsky et al., 1997) and (c.f. (Rothenfluh et al., 2000a, 2000b)), we expected a phase difference between TUG-activated LABL luminescence signal and PER-BG::Luc signal. As anticipated, PER-BG::Luc luminescence signal was phase-delayed compared to both TUG-activated LABL and *plo* flies. Thus, LABL was predictably comparable to both *plo* and PER-BG::Luc flies.

**Figure 3.**
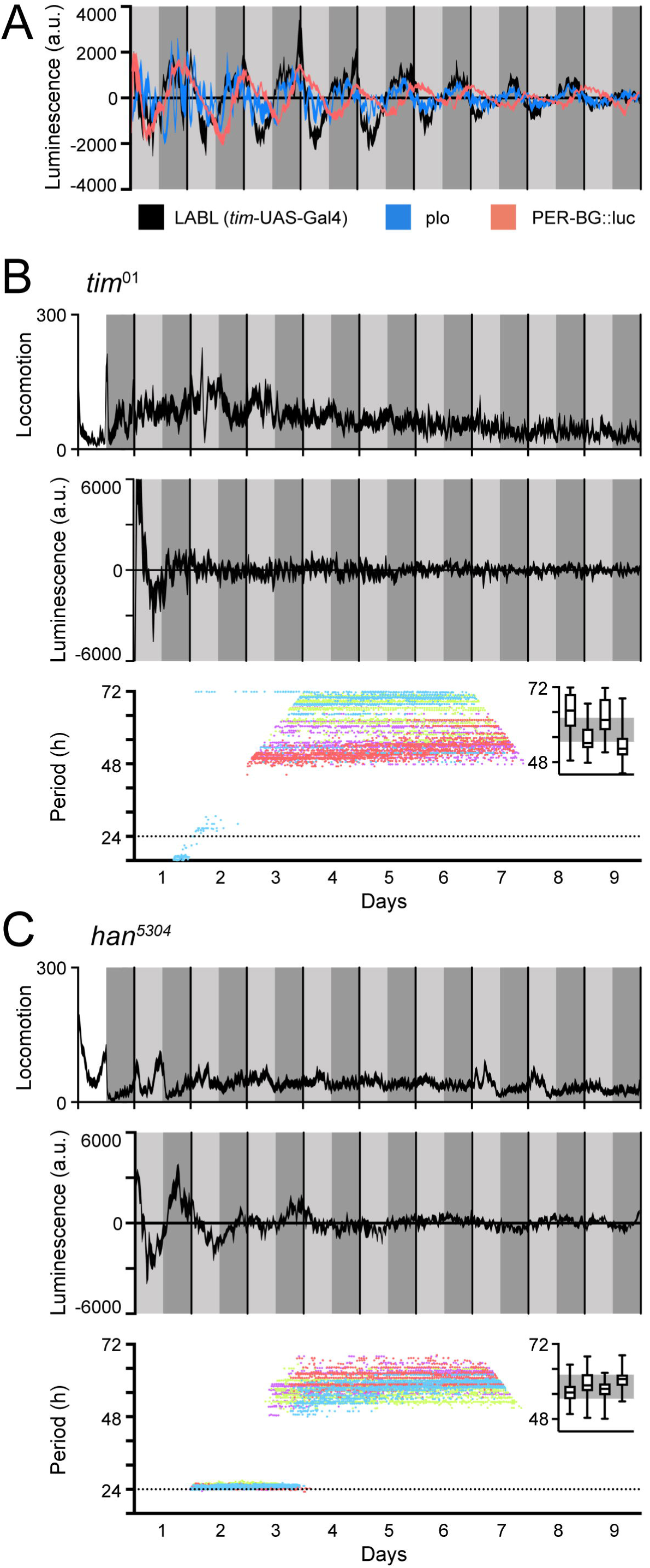
LABL oscillations are comparable to other luminescence reporters and respond to circadian clock manipulation. (**A**) Luminescence reporters compared. LABL activated by *tim*-UAS-Gal4 (black curve) is compared to per promoter fused to luciferase (plo; blue curve) and PER-BG::Luc (pink curve). (**B**) LABL is activated by *tim*-UAS-Gal4 in a *tim*^01^ genetic background. Top graph represents locomotor activity. Locomotion activity is measured in beam breaks (counts) per 30-minute bouts. Curve represents rhythmic locomotor activity of 25 flies, +/- SEM (thickness of curve). White background indicates lights on. Vertical solid lines divide days. Light grey and dark grey backgrounds represent subjective day and subjective night, respectively. Middle graph illustrates luminescence signal over time. Luminescence signal is normalized to exponential decay of signal, values averaged into 30-minute bins, as the mean of four experiments presented, +/- SEM (thickness of curve). Light grey and dark grey backgrounds represent subjective day and subjective night, respectively. Vertical solid black lines divide days. Lower graph illustrates changes in oscillation period over time determined by Morlet wavelet fitting. Parameters are as described in Figure 2E. Each color dots represent an experimental replicate of wavelet period best-fits in the *tim*^01^ genetic background. Inset is a box plot of period best-fits of each of the four experimental replicates, plotted. Grey background represents interquartile range of box plots of the *han*^5304^ genetic background. (**C**) LABL is activated by *tim*-UAS-Gal4 in a *han*^5304^ genetic background. Graphs are as described in panel B.

Eliminating the clock causes behavioural arrhythmicity and loss of transcriptional oscillation. To ensure that the transcription oscillations we observed using LABL were clock-dependent, we monitored TUG-activated LABL luminescence oscillations in a genetic background lacking *tim* expression (*tim*^01^) (Figure 3B). As expected, both locomotor activity and transcription oscillation of *tim*^01^ flies were arrhythmic (top and middle panel). Importantly, our attempt to fit Morlet wavelets to the luminescence data revealed inconsistent quantifications of period, consistent with arrhythmic oscillation. Elimination of PDF signaling using *han*^5304^ mutant flies (expressing PDF receptor lacking a cytoplasmic tail) permitted rhythmic behavior in the first 1-2 days of constant darkness, but ultimately resulted in arrhythmic behavior (cf. (Hyun et al., 2005)). When we characterized LABL oscillations in a *han*^5304^ background (Figure 3C), we found that flies became arrhythmic in their locomotor activity in the second day of constant darkness, as expected. While there was also a rapid decay in transcription oscillation, measured by LABL, closer visual inspection revealed that ∼24 h oscillations decayed into ∼2-3-day infradian oscillations. Quantification of this period across time revealed a stable, reproducible oscillation with a ∼60 h period. Such a coherent 60-hour infradian oscillation of luminescence was not observable in flies lacking a functional clock, suggesting that these infradian oscillations are clock-dependent. While rhythmic and clock-dependent, we interpret these infradian oscillations to represent arrythmia from the 24-hour circadian perspective (i.e., they oscillate with a period on the order of days). These data together demonstrated that the LABL signal was clock-dependent, PDF receptor (PDFR)-dependent, and that it reflected circadian locomotor behavior. Importantly, our method of analysis quantified mutation-caused changes in transcription oscillations robustly and consistently.

### Anatomical expression and LABL activation of some commonly used Gal4 lines

Although the anatomical locations targeted by circadian Gal4 drivers are well characterized in the brain (Figure 4A), it is possible that these driver lines express Gal4 in non-neuronal tissue or non-circadian neurons as well. To establish the anatomical regions in which different Gal4 lines activate LABL, we tested luminescence in whole animals, distinguishing between Gal4 lines that visibly activate LABL in the body from those which do not. Flies in which LABL was activated using Clk4.1-Gal4, Mai179-Gal4 and Clk9M-Gal4 exhibited luminescence signal from the body, similar to that of TUG (Figure 4B). On the other hand, the Pdf-, DvPdf- and R18H11-Gal4 drivers showed no luminescence in peripheral tissues (Figure 4B). We therefore monitored luminescence from explanted live fly brains from Pdf-, DvPdf- and R18H11-Gal4-activated LABL flies (Figure 4C). We found that Pdf-Gal4 activated LABL flies emitted luminescence from a single point located in the region where the LNvs are expected to be. DvPdf-Gal4 activated LABL flies emitted luminescence from a wider region, suggesting that the sources of light are the LNvs and LNds. R18H11-Gal4 activated LABL fly brains emitted luminescence from the dorsal area, in the region DN1s are located. These observations correlated well with expected expression pattern of these Gal4 lines (Figure 4A). To ensure that the luminescence signal recorded from the brain specifically reflected the circadian neuronal cluster in which the Gal4 drivers were expected to activate LABL, we tested for FLP activity using G-TRACE reporter flies co-stained for TIM expression (Figure 4D). G-TRACE uses the Ubip63 promoter fused to stop codons flanked by FRT sequences that are in turn fused to GFP (Evans et al., 2009). Red-stinger (RFP)-positive neurons revealed Gal4-active sites, as shown. GFP-positive neurons that co-stain with TIM antibody revealed that the LNvs (Pdf-Gal4), the LNvs, 5th s-LNv and 3 LNd neurons (DvPdf-Gal4), and DN1s (R18H11-Gal4) had the expected FLP activity. Thus, Pdf-, DvPdf- and R18H11-Gal4 lines activated LABL in the predicted neuronal clusters, indicating that they are suitable for use with LABL to monitor neuron-specific clocks.

**Figure 4.**
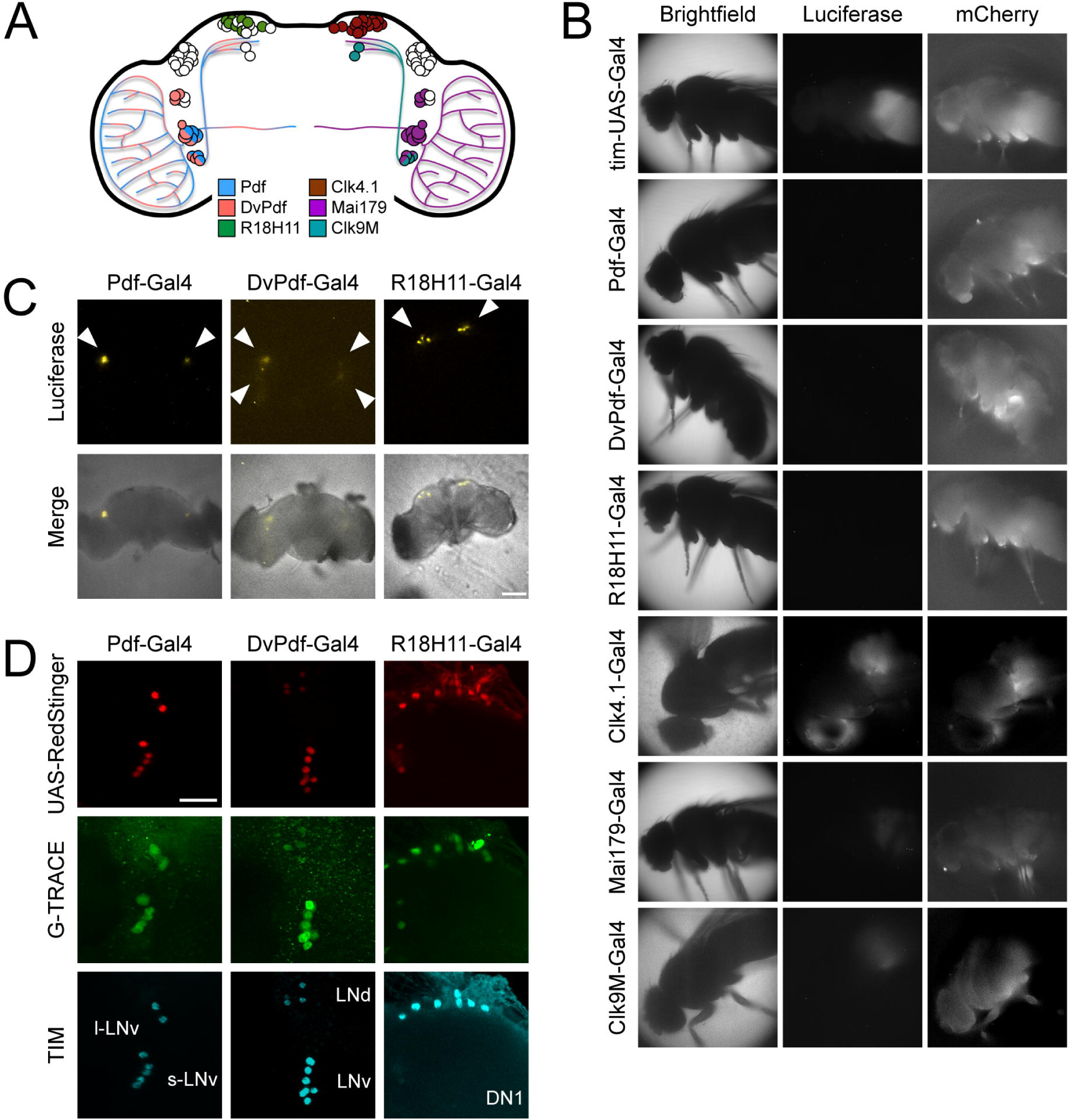
Different circadian Gal4 drivers activate LABL in distinct neurons and tissues. (**A**) Schematic of a *Drosophila* brain and Gal4 driver expression patterns. Pdf-Gal4 (blue): LNvs. DvPdf-Gal4 (pink) and Mai179 (purple): LNvs, 3 LNds and 5th s-LNv. R18H11-Gal4 (green) and Clk4.1 (brown): DN1s. Clk9M (teal): DN2s and small LNvs. (**B**) Luminescence and fluorescence images of whole flies. Flies in bright field (first column), imaged for luminescence (Luciferase; second column) and fluorescence (mCherry; third column). Drivers described in panel A were used to activate LABL in the flies, as indicated. (**C**) Luminescence images of explanted brains. Explanted live brains were imaged for luminescence (Luciferase; top row) and imaged in bright field merged with luminescence (bottom row) to determine sites of LABL activation by the indicated Gal4 drivers. White arrow heads denote estimated location of clock neurons: LNvs in Pdf-Gal4 neurons, LNvs and LNds in DvPdf-Gal4 neurons and DN1s in R18H11-Gal4 neurons. Scale bar represents 50 μm. (**D**) Sites of Flipase activity in fly brains. Fly lines expressing UAS-RedStinger (red fluorescence protein, shown in red) and the indicated driver were crossed into G-TRACE flies and brains were stained for TIM (cyan), RedStinger (red) and GFP (G-TRACE; green). Identity of fluorescence neurons labelled in bottom row. l-LNv: large ventral lateral neurons. s-LNv: small ventral lateral neurons. LNd: dorsal lateral neurons. DN1: dorsal neurons 1. GFP expression is indicative of FLP activity. Pdf-Gal4 driver targets LNvs, DvPdf-Gal4 driver targets LNvs, 5th s-LNv, 3 LNds, and R18H11-Gal4 driver targets DN1s. Scale bar represents 20 μm.

### Elimination of PDF signaling has variable effects on neuronal clocks

Elimination of the clock component *tim* causes arrhythmic locomotor activity and clock oscillations as measured by LABL (Figure 3B). Elimination of PDF signaling leads to arrhythmic locomotor activity in constant darkness and infradian clock oscillations (Figure 3C), which may be the result of non-synchronized clocks within the fly (Lin et al., 2004; Yoshii et al., 2009). We therefore sought to directly measure clock activity in distinct circadian neurons through activation of LABL in mutants lacking PDF signaling (*han*^5304^). Elimination of PDF signaling revealed two categories of Gal4 lines: those which continued to oscillate and those which did not (Figure 5A). Pdf-, DvPdf-, and R18H11-Gal4 drivers (collectively, “brain drivers”) retained transcription oscillation in the absence of PDF signaling for a few days, while Clk4.1-, Mai179-, Clk9M-Gal4, and TUG drivers (collectively, “body drivers”) began to deteriorate after one cycle of oscillation. This deterioration could be observed in the earlier stability constants (see vertical dashed lines) of flies with a *han*^5304^ genetic background (colored curves) compared to wild type flies (black curves). This result suggests that the non-neuronal circadian tissues in which the body drivers are expressed rely on PDFR to maintain their circadian clocks.

**Figure 5.**
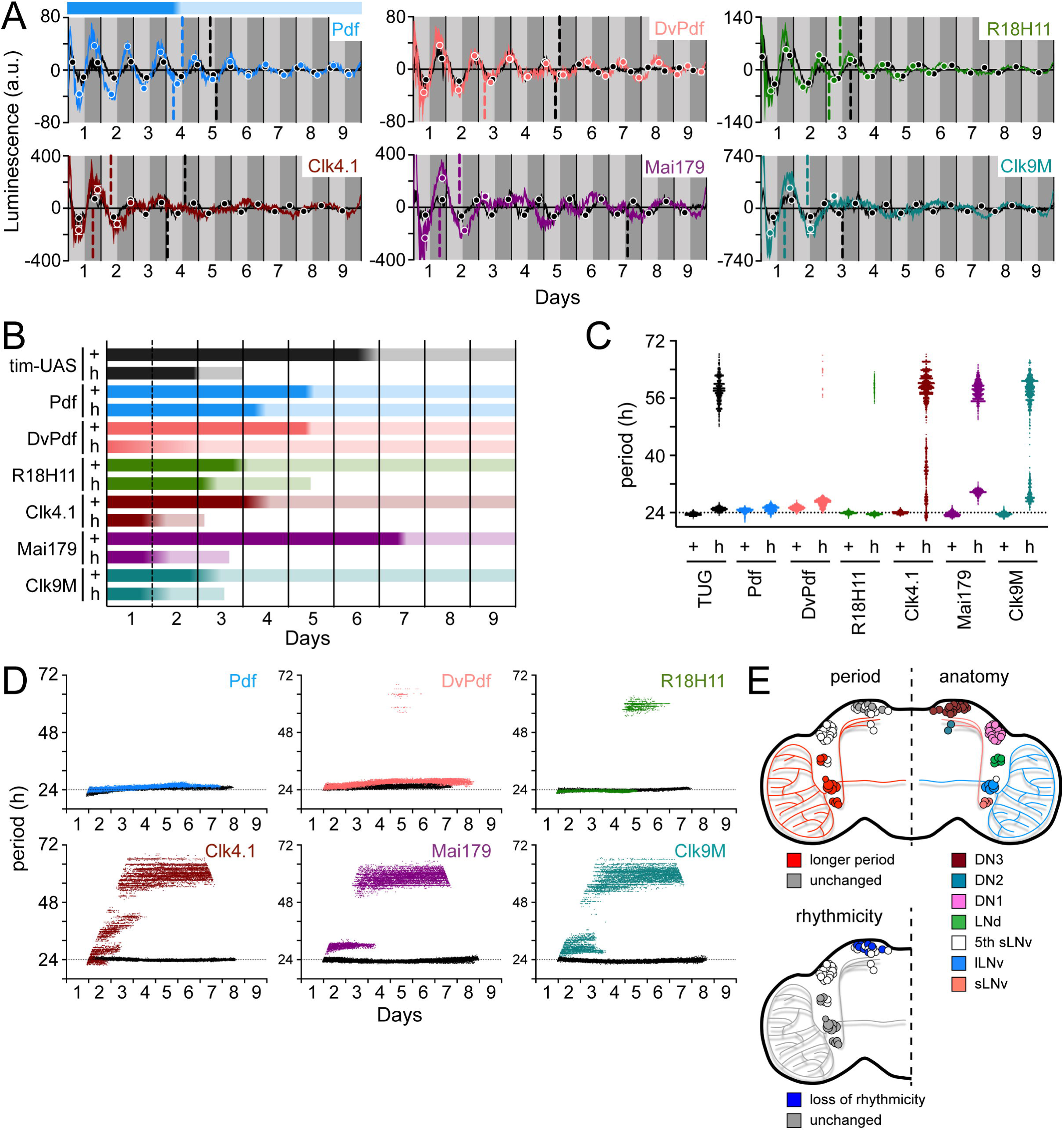
Different circadian Gal4 drivers create distinct oscillation patterns that respond to loss of PDF signaling differently. (**A**) Luminescence oscillations measured from LABL flies activated by the indicated Gal4 drivers, in a wild type or PDF signaling deficient genetic background, plotted over time. Colored curves represent signal from flies with no PDF signaling (*han*^5304^) compared to their wild type counterparts (black curves). Drivers used are Pdf- (light blue), DvPdf- (pink), R18H11- (green), Clk4.1- (brown), Mai179- (purple) and Clk9M-Gal4 (cyan). Otherwise, graphs are as described in Figure 2C. Bar above the curve is a visual representation of the amplitude decay across time: color fade represents the amplitude stability constant. Example given is the decay of Pdf-activated LABL in a *han*^5304^ genetic background, in blue. (**B**) Stability of luminescence oscillations from LABL flies expressing different drivers are quantified in a wild type (+) and PDF signaling-deficient (*han*^5304^; h) background. Colors correlate with Gal4 drivers described in panel A. Color fade in the bars represent the location of the amplitude stability constant (point of inflections of the decay in amplitude of oscillation). Length of bar indicates time at which luminescence oscillation becomes arrhythmic. (**C**) Differences in average period of luminescence oscillation measured by Morlet wavelet fitting, comparing wild type (+) and PDF signaling deficient (*han*^5304^; h) flies. Colors correlate with Gal4 drivers described in panel A. If arrhythmic (>48 h) period fits are excluded, periods of oscillations measured in the brain for wild type and *han*^5304^ flies can be averaged, +/- SD. Wild type: Pdf, 24.53 +/- 0.61; DvPdf, 25.33 +/- 0.58; R18H11, 23.93 +/- 0.33 hours. *han*^5304^: Pdf, 25.08 +/- 0.78; DvPdf, 26.79 +/- 1.02; R18H11, 23.58 +/- 0.30 hours. The dotted horizontal line denotes 24 hours as a point of reference. (**D**) Changes in oscillation period over time determined by Morlet wavelet fitting. Colored dots represent best fits from flies with no PDF signaling (*han*^5304^) compared to their wild type counterparts (black curves). Colors correlate with Gal4 drivers described in panel A. The dotted horizontal line denotes 24 hours as a point of reference. (**E**) Schematic summary of changes in period and rhythmicity of luminescence oscillation in distinct parts of the fly brain caused by loss of PDF signaling (*han*^5304^ genetic background). LNvs and 3 LNds exhibit longer luminescence oscillation period (top left hemisphere, red) but remain rhythmic (bottom left hemisphere, grey). Luminescence oscillation in DN1s maintain the same period with loss of PDF signaling (top left hemisphere, grey), but become arrhythmic over time (bottom left hemisphere, blue). Anatomical location of s-LNvs (small ventral lateral neurons), l-LNvs (large ventral lateral neurons), 5th s-LNv (5th small ventral lateral neuron), LNds (dorsal lateral neurons), and DN1s, DN2s, DN3s (dorsal neurons 1, 2, 3), as shown (top right hemisphere).

To identify differential tissue-specific clock stability, we examined the stability of different neuron-specific clocks in both wild-type controls and PDFR mutants. We found that neuron-specific clocks demonstrated differential dependence on PDFR for circadian oscillation of transcription in stability, period, and amplitude. In a wild-type genetic background, the stability of luminescence oscillation depended on where LABL has been activated. Here we represent the change in amplitude as a bar (see top of luminescence recordings, Figure 5A), with the dark region representing the early minima/maxima within the S-curve, the light region representing the later minima/maxima within the S-curve, and the gradient representing the points of inflection observed. A comparison of clock amplitude stability revealed that Mai179-activated wild type flies exhibited an amplitude stability constant (defined in Figure 2C) of 7 days in constant darkness, while Clk9M-activated wild type flies exhibited a stability constant of 2 days (Figure 5B). In all cases, elimination of functional PDFR shifted the stability constant earlier in constant dark conditions. All LABL oscillations activated by body drivers fell into arrhythmia by the end of Day 3 in constant darkness, which is consistent with arrhythmic behaviour observed in locomotor activity (Figure 3C). Interestingly, with loss of functional PDFR, R18H11-activated LABL flies had a relatively unchanged (∼ half day difference) stability constant but rely on PDF signaling for maintaining rhythmicity (arrhythmic after Day 5). Pdf- and DvPdf-activated *han*^5304^ LABL flies maintained their oscillations with some instability (Figure 5B), in agreement with earlier reports involving LNv and LNd oscillations (Lin et al., 2004). The advanced stability constant in DvPDF-activated *han*^5304^ LABL flies is possibly due to phase dispersal in the s-LNvs and phase advance in the LNds caused by a loss of PDF signaling (Lin et al., 2004). Thus, body-driver activated LABL flies are dependent on functional PDFR to maintain rhythmic oscillations. Of the brain drivers, R18H11-activated LABL flies were dependent on functional PDFR to maintain rhythmic oscillations in the long term (beyond 5 days), but Pdf- and DvPdf-activated LABL flies continued to oscillate in flies lacking PDF signaling. Thus, PDF signaling is not necessary to maintain lateral neuron clock oscillations, as also inferred by others in *pdf*^01^ flies (Yoshii et al., 2009).

Flies lacking PDF (*pdf*^01^) exhibit shorter behavioural period as well as advanced PER nuclear translocation in LNds (Lin et al., 2004; Renn et al., 1999). We therefore quantified changes in period of transcription oscillations in the different clocks to determine if shortened behavioural rhythms were caused by shortened clock oscillatory periods. Examination of the averaged peaks and troughs of transcription oscillation (Figure 5A) revealed a consistent phase advance in flies lacking functional PDFR (despite longer oscillation periods), which may partially account for the observed advance in PER nuclear translocation in LNds. Surprisingly, elimination of functional PDFR caused longer observed clock oscillation periods in different regions of the brain, except those where R18H11-Gal4 was used to activate LABL flies, which maintained wild-type-like oscillatory periods (Figure 5C). This is inconsistent with previously published locomotor data that shows a shortened period of *pdf*^01^ flies, when arrhythmic animals are excluded (Renn et al., 1999). Therefore, the observed shorter behavioural rhythms may be caused by other clocks that oscillate with a shorter period, or elimination of PDFR may not entirely reflect the effect of a *pdf*^01^ fly. Indeed, PDFR may be responsive to other neuropeptides (e.g. DH31) (Mertens et al., 2005).

Periods of luminescence oscillation of different tissues in *han*^5304^ flies were lengthened to different degrees. Luminescence from Mai179-activated LABL in *han*^5304^ flies exhibited a ∼30-hour period before becoming arrhythmic on day 3, as compared to a ∼24-hour period in wild type flies (Figure 5C). A time course revealed that all oscillators that lose rhythmicity due to loss of PDF signaling appeared to form stable ∼60-hour infradian rhythms (Figure 5D). Of these, the body drivers (TUG, Clk4.1-, Mai179-, Clk9M-Gal4) appeared to show a more disorganized period decay in constant darkness (Figure 5D). Pdf-activated LABL flies exhibited a higher amplitude for a longer amount of time (∼5 days) in the absence of PDF signaling as compared to the first cycle of oscillation. This is similar to an observed increase in *tim* promoter activity in s-LNvs in flies lacking PDF signaling (Mezan et al., 2016). On the other hand, DvPdf- and R18H11-activated *han*^5304^ LABL flies exhibited amplitude peaks similar to their wild-type counterparts after the first cycle of oscillation. This rapid loss of difference in amplitude after the first day was also observed in body-driver activated LABL flies. From these data, we confirm that PDF signaling is critical for rhythmic locomotor activity and some neuronal clocks but is not necessary to sustain the “master clock” neurons (LNvs) (Lin et al., 2004; Yoshii et al., 2009). Importantly, we conclude from these data that LABL can demonstrate that PDF signaling has distinct effects on different neurons and tissues in clock stability, period, and amplitude (Figure 5E).

### Peripheral clocks exhibit distinct oscillations and variable PDF-dependence

Given the differences in clock transcription oscillation in LABL flies activated by body-drivers and brain-drivers, we decided to directly measure clock oscillations in specific peripheral organs. To characterize tissue-specific peripheral clock drivers, we monitored the stability of tissue-specific clock function in constant dark conditions in both wild-type controls and *pdfr* mutants. Specifically, we examined LABL oscillations using the neuronal driver elav-Gal4 (Saitoe et al., 2001; Schuster et al., 1996; Sink et al., 2001), intestinal drivers esg- and NP3084-Gal4 (Bonnay et al., 2013; Croker et al., 2007; Lee et al., 2016; Strand and Micchelli, 2011), fat body drivers C564- and LSP2-Gal4 (C564 also expresses in hemocytes and some male reproductive tissues) (Hrdlicka et al., 2002; Paredes et al., 2011; Takeuchi et al., 2015; Zaidman-Rémy et al., 2006), and muscle driver mef2 (also expresses in some neurons [LNvs]) (Blanchard et al., 2010; Cripps et al., 1998; Gajewski et al., 1997) (Figure 6A). In flies both with and lacking PDF signaling, all drivers (except NP3084-Gal4) activated LABL to reveal consistent transcription oscillations (Figure 6B). Interestingly, mef2- and esg-driven LABL flies lacking PDF signaling exhibited a loss of measurable oscillation between days 5-6 and 4-6, respectively, followed by a re-established rhythm. Quantification of amplitude stability revealed that LABL driven by these drivers exclusively (mef2- and esg-Gal4) exhibited earlier stability constants in flies lacking PDF signaling, compared to their wild type counterparts (Figure 6C). The equivalent stability constants of peripheral clocks between flies with and lacking PDF signaling suggests that PDF signaling is not critical in maintaining these oscillators.

**Figure 6.**
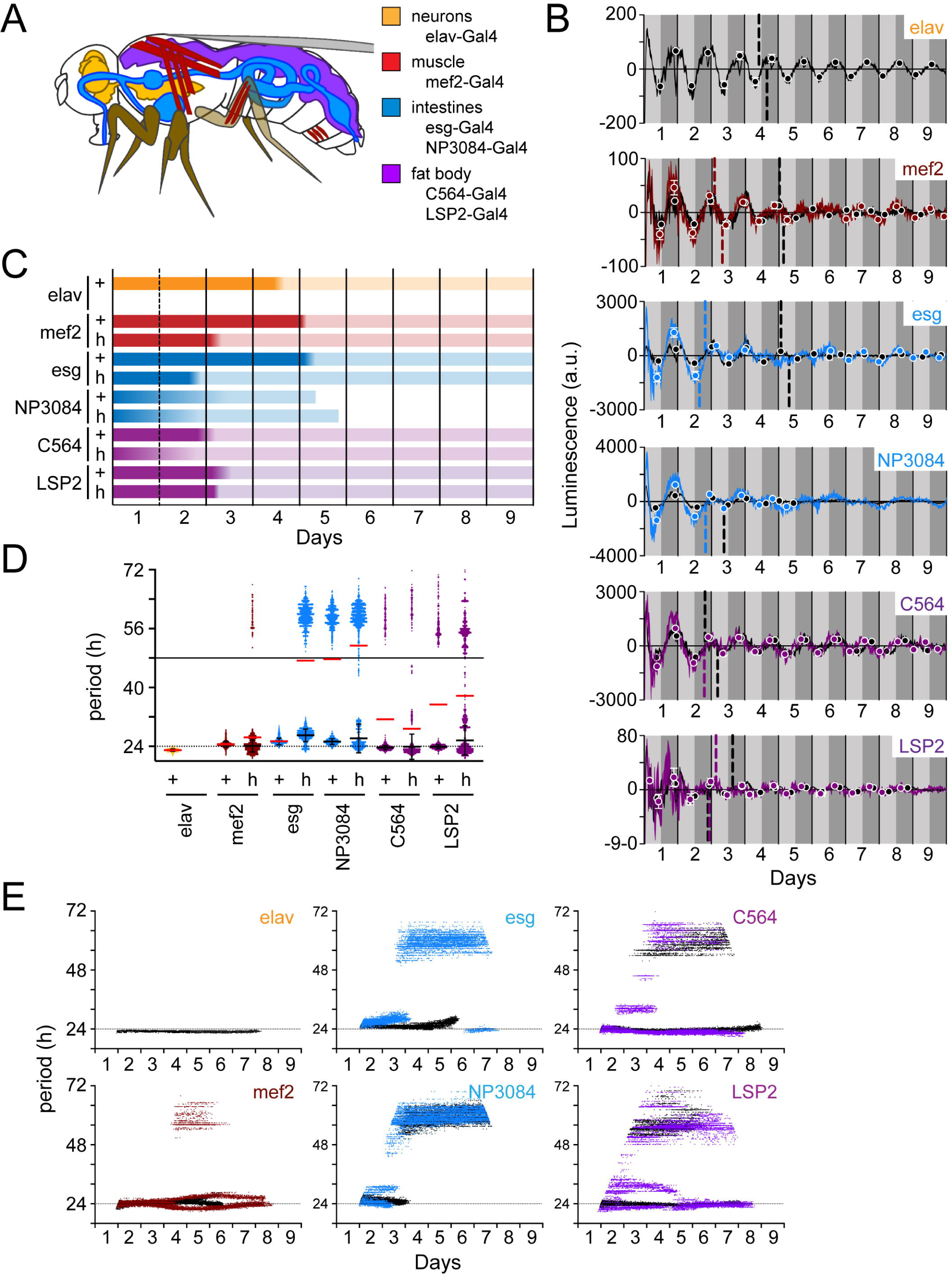
Peripheral clocks have distinct characteristics in a wild type and PDF receptor mutant genetic background. (**A**) Schematic of target expression pattern of peripheral tissue Gal4 drivers in *Drosophila*. Elav- (neurons; orange), mef2- (muscle; red), esg- and NP3084- (intestines; blue), and C564- and LSP2-Gal4 (fat body; purple). Note that C564-Gal4 also expresses in hemocytes and some male reproductive tissues and mef2-Gal4 has been reported to express LNv neurons. (**B**) Luminescence oscillations measured from LABL flies activated by the indicated Gal4 drivers in peripheral tissues, in a wild type or PDF signaling deficient genetic background, plotted over time. Colored curves represent signal from flies with no PDF signaling (*han*^5304^) compared to their wild type counterparts (black curves). Drivers used are as described in panel A. Graphs are as described in Figure 2C. (**C**) Stability of luminescence oscillations from LABL flies expressing different drivers are quantified in a wild type (+) and PDF signaling deficient (*han*^5304^; h) background. Colors correlate with Gal4 drivers described in panel A. Color fade in the bars represent the location of the amplitude stability constant (point of inflections of the decay in amplitude of oscillation). Length of bar indicates time at which luminescence oscillation becomes arrhythmic. (**D**) Differences in average period of luminescence oscillation measured by Morlet wavelet fitting, comparing wild type (+) and PDF signaling deficient (*han*^5304^; h) flies. Colors correlate with Gal4 drivers described in panel A. Black bars represent mean period of “rhythmic” oscillations early in measurement, +/-SEM. Red bars represent mean period of all wavelet fits (including those >48 h). The dotted and solid horizontal lines denote 24 and 48 hours as a point of reference, respectively. (**E**) Changes in oscillation period over time determined by Morlet wavelet fitting. Colored dots represent best fits from flies with no PDF signaling (*han*^5304^) compared to their wild type counterparts (black curves). Colors correlate with Gal4 drivers described in panel A. The dotted horizontal line denotes 24 hours as a point of reference.

Quantification of clock periods in peripheral tissues revealed distinct periods in wild type flies (Figure 6D). Importantly, LABL activated by fat body-specific drivers (C564 and LSP2) exhibited oscillation periods greater than 48 hours, suggesting that these clocks may not be as robust (e.g. compared to elav+ clocks) in constant darkness. Elimination of PDF signaling appeared to destabilize the period of oscillation of all measured clocks, to varying degrees (Figure 6D). To uncover the extent of period stability in the absence of PDF signaling, we monitored changes in period of peripheral clocks over time (Figure 6E). Intestine-specific Gal4 drivers revealed transcriptional oscillations that stabilized in an infradian ∼60-hour period in flies lacking PDF signaling, similar to that observed using “body clock drivers (Figure 5D). On the other hand, muscle- and fat body-specific drivers appeared to maintain a rhythmic ∼24- hour transcription oscillation, though these were unstable, as demonstrated by intermittent deviations (Figure 6E). We concluded from these data that PDF signaling may be required to stabilize an oscillator, but not to sustain it. Interestingly, periods of clock oscillation in the fat body were shorter in the absence of PDF signaling as compared to wild type (Figures 6B and 6D), contrasting the longer periods observed in the same mutant background in Pdf- and DvPdf-activated LABL flies (Figure 5C), but similar to shorter behavioural periods measured by locomotion (Hyun et al., 2005). Thus, the LABL reporter system was able to distinguish between loss of rhythmic oscillation and destabilized oscillation, as well as changes in the period of transcription oscillation, in peripheral tissue.

## DISCUSSION

Understanding the links among circadian rhythms, behavioral/metabolic disorders, and the hierarchical organization of circadian clocks requires measurement of these distinct clocks. Here, we present a method in which we expressed Luciferase under the control of a circadian promoter in targeted cells or tissues, allowing us to directly quantify the oscillatory properties of distinct clocks. Using this method, we found that clocks in different tissues oscillated differently, underscoring two important features of Drosophila circadian clocks: (1) circadian clocks have distinct oscillatory properties, suggesting that they are differently regulated; and (2) the PDF+ LNvs do not exert complete dominance over other clocks through PDF signaling.

Distinct neuronal populations that express circadian clocks have different roles in regulating circadian behavior. Previous work by others analyzing locomotion patterns found that LNvs regulate morning anticipatory behaviour and rhythmic locomotion in constant darkness, and LNds (and the 5th sLNv) regulate evening anticipation and rhythmic locomotion in constant light conditions depending on the genetic background (Murad et al., 2007; Picot et al., 2007). These different responsibilities for distinct neurons suggest that coherent behavioral rhythms are the result of converged function of the entire circadian neuronal network. It follows that distinct neuronal functions may necessitate the need for different molecular mechanisms to regulate individual clocks. Indeed, Casein Kinase II, a regulator of PER/TIM repressor complex nuclear accumulation, is expressed in the LNvs but not in other circadian neurons (Top et al., 2016). LABL offers an opportunity to explore the effect of mutations and different environmental factors on these distinct clocks.

LABL offers several advantages in its use as a reporter. Unlike other systems that rely on measuring luminescence signal from explanted fly brains, LABL leaves the brain and other tissues in place, allowing them to remain responsive to physiological input from the body of the animal. Current techniques that remove tissues from the context of the whole body measure clock oscillations in the absence of physiological regulatory factors. For example, explanted brains reveal ∼29 h tim-luc oscillations and ∼25 h 8.0-luc oscillations (a *per* fragment that expresses PER::luciferase fusion protein, while missing a majority of its transcription regulatory sequence) (Versteven et al., 2020), and differences in l-LNv neuronal firing are explained by the presence of eyes (Muraro and Ceriani, 2015), suggesting that a number of external factors maintain physiological oscillations. LABL also allows animals to roam freely, avoiding any aberrations that may arise from tethering the head of the animal to an imaging stage. This technique does rely on the specific expression of Gal4 in the intended target tissues. For example, in characterizing some common circadian Gal4 lines, such as Clk4.1, Mai179 and Clk9M, we found that they expressed Gal4 in peripheral tissues, which disqualifies use of these drivers for neuron-specific LABL-mediated monitoring of transcriptional oscillations (Figure 4). LABL is distinct from other *in vivo* luminescence-based reporters. Reporters such as *plo* and PER-BG::Luc report the cumulative oscillation of either the *period* promoter or Period protein in the whole animal (Figure 3A). Other reporters rely on the oscillation of calcium levels (Guo et al., 2017) or the oscillation of secondary signaling such as NFκB or CREB2 (Tanenhaus et al., 2012), which measure cell responses to extracellular signals, but not the circadian clock itself. While these methods are effective in measuring how targeted tissues responds to signals from elsewhere, they do not directly measure the transcription/translation feedback loop. In contrast, LABL permits the direct measurement of the circadian clock in a target cell or target tissue, which is critical to understanding the effect of a mutation or environmental input on distinct clocks, particularly non-neuronal or peripheral clocks that may not use the same signaling outputs as neuronal clocks.

Using LABL, we confirmed that distinct circadian clocks exhibit distinct characteristics. We noted, for example, a sharp dip in luminescence that bisects the first peak of oscillation in all measured flies (Figure 2C, Supplementary Figure 2), except with PER-BG::Luc flies (Figure 3A), suggesting that this may be a transcription regulation-specific phenomenon, as previously suggested by others (Brandes et al., 1996). We also observed that not all oscillators are maintained under constant conditions (e.g., NP3084-activated LABL) (Figure 6), suggesting that a subset of clocks (i.e. the transcription feedback loop) may at times be sustained by environmental or other forms of input. Such differences in sustaining clock oscillations have been previously observed by others (Roberts et al., 2015; Veleri et al., 2003; Veleri and Wülbeck, 2004). For example, the oscillations of the clock protein PER are rapidly dampened in l-LNvs and DN1s in constant dark conditions (compared to other neuronal clusters), while their *per* transcript oscillations are reportedly relatively unaffected, highlighting differences in PER stability versus transcription activity (Roberts et al., 2015; Shafer et al., 2002; Veleri and Wülbeck, 2004; Yang and Sehgal, 2001). We too observed oscillations sustained for days in R18H11-activated LABL flies, i.e. the DN1s (Figures 4 and 5). Thus, LABL allowed similar observations to be made rapidly, without the need for labor-intensive, low time-resolution immunofluorescence staining or *ex vivo* luminescence imaging (Versteven et al., 2020).

Using LABL, we were able to rapidly assess the effects of loss of PDF signaling on tissue-specific clocks as a proof of principle for this method in different mutant or disease contexts. We found that transcriptional oscillations became longer to varying degrees when circadian Gal4 lines are used to activate LABL in flies lacking PDF signaling, or shorter in the case of the fat body clocks. R18H11-activated LABL flies exhibited no period change of transcription oscillation. Oscillations in peripheral tissue became less stable, while remaining intact. In two Gal4 lines, mef2- and esg-Gal4, circadian oscillations were lost through days 5-6, before they were restored later in the time course, perhaps re-established through signals received from other clocks. Finally, the formation of ∼60-hour infradian oscillations in tissues lacking PDF signaling were clock-dependent, since they were distinct from measurement from flies lacking a functional clock, which may be the result of multiple uncoordinated oscillators. It is unlikely that this infradian oscillation was caused by non-synchronised individual flies, since clock oscillations revealed by Pdf-Gal4 persisted despite lacking PDF-signaling, while oscillations revealed by R18H11-Gal4 reverted to infradian rhythms on Day 5. Such infradian rhythms would likely have been missed using classical methods such as immunofluorescence, given their long period. Although describing these infradian oscillations goes beyond the scope of our study, uncovering the nature of these infradian oscillations promises to reveal exciting new features of how distinct clocks are integrated.

The LABL reporter system promises to uncover novel clock mechanisms at increased cellular and temporal resolution. Here, we present data that demonstrate the function and use of this system. LABL has the advantage that it reveals clock oscillations directly, at the intersection of tissues (and cells) in which the *per* promoter is active (i.e. in clock cells) and where Gal4 is expressed, such that cells that express Gal4 but no clock would not interfere with luminescence measurements. LABL reveals distinct clocks that are unique in their character and response to molecular input (e.g. PDF signaling). Together, LABL promises to be an important tool in directly measuring and quantifying subtle changes to individual circadian neuronal sub-types as well as peripheral clocks in *Drosophila*, *in vivo*, in real time, and is likely to find application in other model organisms.

## METHODS

### Fly strains

The following lines were used in the study: tim-UAS-Gal4 (A3) (Blau and Young, 1999), Pdf-Gal4 (Park et al., 2000), DvPdf-Gal4 (Guo et al., 2014), R18H11-Gal4 (Guo et al., 2016), Clk4.1-Gal4 (L. Zhang et al., 2010; Y. Zhang et al., 2010) (gift from Michael W. Young); Mai179-Gal4 (Picot et al., 2007) (gift from C. Helfrich-Foerster); Clk9M-Gal4 (Kaneko et al., 2012) (gift from O. T. Shafer); 3xUAS-FLP2::pest (Nern et al., 2011) (Janelia Research Campus); PDF receptor mutant *han*^5304^ (Hyun et al., 2005; Mertens et al., 2005) (gift from P. H. Taghert). G-TRACE flies were acquired from BDSC (32251) (Evans et al., 2009). Genotypes of all flies used in experiments are summarized in Supplementary Table 1. Flies were reared on standard cornmeal/agar/yeast/molasses medium at room temperature (22 °C) or 18 °C in ambient laboratory light. Flies were entrained in a light-dark cycle (12h:12h) at 25 °C.

### LABL flies

LABL was constructed into an attB cloning vector for Drosophila embryo injection and PhiC31-mediated genome integration (Bischof et al., 2007) (Supplementary Figure 1). The minimal *per* promoter (Bargiello et al., 1984) (ranging from restriction sites SphI to XbaI) was first cloned into the pattB vector. The 5’ untranslated region, ranging from XbaI site to the *per* ATG start codon was rebuilt using standard PCR: the ATG was conserved and an FRT sequence followed by a NotI restriction site was added. mCherry was amplified using standard PCR, its start codon eliminated, and a 5’ NotI restriction site, three 3’ stop codons and an EcoRI restriction site added. Luc2 (Promega) was amplified using standard PCR its start codon eliminated, a 5’ EcoRI restriction site followed by FRT sequence added and then 3’ XhoI and KpnI restriction sites added. The plasmid was then assembled and injected into embryos (BestGene Inc.).

### Luminescence assay

Luminescence of flies were measured using the LumiCycle 32 Color (Actimetrics). Custom 35 mm plates (designed by Actimetrics) were used to adapt the LumiCycle 32 for *Drosophila* use. D-luciferin potassium salt (Cayman Chemicals or Gold Biotechnology) was mixed with standard fly food to a final concentration of 15 mM. Volume of food in the plate was sufficient to limit fly movement along the z-axis of the plates. 15 flies were placed in each plate and covered with a cover slip. Each of the four replicates were recorded in a different position in the luminometer to ensure equal representation of data from each of the four photomultiplier tubes. Luminescence from each plate was recorded in 4-minute intervals on a standard Windows-operated PC using software by Actimetrics.

Luminescence of transfected S2 cells were measured using a liquid scintillation counter LS6000IC (Beckman) in single-photon collection mode. Cells were transfected in 6-well plates at 80% confluency using Effectene transfection reagent (Qiagen) following the manufacturer’s protocol. Cells were transfected with 200 ng total DNA with the indicated plasmids distributed in equivalent amounts. Plasmids used were pAc-*clk*-CFP (actin promoter driven *clock* fused to CFP) (Top et al., 2018), pAc-*flp* (actin driven *flipase*; gift from Nicholas Stavropoulos), LABL plasmid (above). Cells were fed 24 h after transfection and lysed another 24 h later in Cell Culture Lysis Reagent (Promega). Extracts were mixed with Luciferase Assay Reagent (Promega) in a 1:5 ratio and monitored for luminescence.

### Luminescence analysis

Actimetrics analysis software was used to normalize the exponential decay of luminescence signal over days using a polynomial curve fit, with no smoothing. Data were exported into .csv files for analysis. Custom python code was used to organize luminescence data into 30-minute bins and quantify peaks, troughs and decay of oscillating luminescence signal (LABLv9.py; www.top-lab.org/downloads). A sinusoidal curve was fitted to each day, +/- half a day, with the x- and y- coordinates for the local maxima and minima identified and quantified. Overlapping values for each peak and trough were averaged for x- and y- coordinates before being plotted, +/- standard deviation. Graphpad Prism 9 software was used to fit an S-curve to these points, and the calculated value for IC50 was used to identify the point of inflection to identify the “stability constant”. Custom python code was also used to quantify periods of oscillations using a Morlet wavelet fit (waveletsv4.py; www.top-lab.org/downloads), as previously described by others (Leise and Harrington, 2011). The bottom 25% of best fits were omitted from analysis. Computer-generated oscillation data was used to confirm the parameters for analysis (data not shown): bandwidth = 3; central frequency = 1; data points in an hour = 15 (i.e., 15 x 4 min in an hour); zero-hour mark = 10 (10:00 am, subjective dawn); period of shortest wave match = 16; period of longest wave match = 72; increment of wavelet fit = 1; percent threshold for discarding data as a fraction = 0.25. Highest confidence interval values were used to plot period across time. The time (x-value) when an oscillation was considered arrhythmic was defined as the last 25 data points before a wavelet fit exhibited a jump from ∼24 hours to ∼60 hours, and averaged. All four replicates that showed a shift to circadian arrhythmicity were averaged and plotted.

### Locomotor activity

Individual flies were analyzed for locomotor activity for three days in a light-dark cycle, followed by seven days in constant darkness using the *Drosophila* Activity Monitor System 5 (Trikinetics). Circadian behaviour was determined in the analysis of time in constant darkness. Periodicity of rhythmic locomotor activity was determined using the ImageJ plug-in ActogramJ (Schmid et al., 2011).

### Fluorescence microscopy and immunohistochemistry

To measure loss of mCherry signal from LABL plasmid in S2 cells, cells were grown in Lab-Tek II chamber slides (Nunc, Rochester, NY). Cells were imaged using a DeltaVision system (Applied Precision, Issaquah, WA) equipped with an inverted Olympus IX70 microscope (60X oil objective, 1.42 N.A.), a CFP/YFP/mCherry filter set and dichroic mirror (Chroma, Foothill Ranch, CA), a CCD camera (Photometrics, Tucson, AZ), and an XYZ piezoelectric stage for locating and revisiting multiple cells.

Fly brains were collected at ZT23, fixed, mounted, and imaged using Leica confocal microscopy as previously described (Top et al., 2018, 2016). Briefly, fly heads were fixed in PBS with 4% paraformaldehyde and 0.5% Triton X-100. Brains were dissected and washed in PBS with 0.5% Triton X-100. Brains were then probed with a 1:1000 dilution of rat anti-TIM (Myers et al., 1996), a 1:1000 dilution of chicken anti-GFP (Aves Labs), and a 1:1000 dilution of rabbit anti-mCherry (Rockland). Washed brains were re-probed using a 1:500 dilution of goat anti-chicken conjugated to Alexa-488, a 1:200 dilution of donkey anti-rat conjugated to Alexa-594, and a 1:200 dilution of donkey anti-rabbit conjugated to Alexa-647 secondary antibodies (Jackson Immunological). Brains were mounted using Fluoromount G (Beckman Coulter, Brea, CA) and imaged using a ZEISS LSM 710 confocal microscopy at 40X magnification.

### Luminescence imaging

Flies were kept at 25°C under light-dark cycles in vials containing 1% agar and 5% sucrose food supplemented with 15mM luciferin for at least 3 days prior to the experiment. Images were taken using a Luminoview LV200 (Olympus) bioluminescence imaging system, equipped with an EM-CCD camera (0.688 MHz EM-CCD CAM-ImageEM X2, Hamamatsu, Japan) at a time point corresponding to the peak phase of expression (between ZT 18 and ZT 20).

Adult flies were anesthetized with ether and dry mounted on a CellView glass bottom imaging dish (35x10mm; Greiner Bio One). Once the position of the fly was determined by bright field microscopy, bioluminescence was measured for 5 minutes with an EM gain of 400, using a UPLSAPO 20x Apochromat, followed by 600ms exposure to fluorescence light. After this, another bright field image was captured and compared to the initial image, in order to ensure that the fly had not moved.

Preparation of the brains was similar to that described previously (Schubert et al., 2020; Versteven et al., 2020). Briefly, brains were dissected in ice cold Ca^2+^ free Ringer’s solution and placed on a CellView glass bottom imaging dish (35x10mm; Greiner Bio One) treated with Heptane glue, where the culture medium was added. The medium contains 20% heat-inactivated fetal bovine serum, 1% Penicillin-Streptomycin and 0.75 µM Luciferin in Schneider’s medium (Sigma-Aldrich). Bioluminescence images were acquired with a UPLSAPO 40x by exposing the brains for 5 minutes with an EM gain of 400. For each brain, a Z-stack of 13 slices was taken, each slice containing a bioluminescence, a fluorescence (200ms), and a bright field image. All images were processed in FIJI (Schindelin et al., 2012) and combined with bright field images in GIMP (https://www.gimp.org/).

## ACKNOWLEDGMENTS

This work was supported by a grant from NSERC (RGPIN-2019-06101) to D.T., from NSF (IOS 1656603) to S.S. and from the Deutsche Forschungsgemeinschaft (DFG) (INST 211/835-1 FUGG, and STA421/7-1) to R.S.

## AUTHOR CONTRIBUTIONS

P.S.J. conceived of and performed experiments, analyzed data. M.O. conceived of and performed experiments, analyzed data. I.T. developed software. S.S. designed computer-generated oscillation data and tested software, assisted in developing software. R.S. analyzed data and wrote manuscript. D.T. conceived of and performed experiments, analyzed data and wrote manuscript.

## COMPETING INTERESTS

None of the authors have any competing interests.

**Supplementary Figure 1. LABL cloning strategy.**

The region of *per* promoter between restriction sites SphI and XbaI (Bargiello et al., 1984) was cloned into pattB cloning vector. The *per* 5’UTR was amplified from plasmid containing the entire *per* minigene using synthetic oligos to introduce an FRT recombination site and NotI restriction site 3’ of the UTR, using standard PCR protocols. The mCherry gene was amplified using standard PCR protocols to introduce a 5’ NotI restriction site and 3’ stop codons and EcoRI restriction site. Luc2 (pGL4.10) was amplified using standard PCR protocols to introduce a 5’ EcoRI restriction site and FRT site and 3’ stop codons and an XhoI restriction site to Luc2. All amplicons were assembled sequentially into the pattB plasmid vector, and sequence confirmed.

**Supplementary Figure 2. Dip in luminescence signal in the first peak of transcription oscillation.**

Luminescence signal recorded from TUG-activated LABL flies presented in Figure 2C was reproduced and the first peak enhanced for clarity. The red box indicates the section of the curve that was enhanced. The red arrow points to the dip in luminescence signal.

## REFERENCES

Acosta-Rodríguez VA, Rijo-Ferreira F, Green CB, Takahashi JS. 2021. Importance of circadian timing for aging and longevity. Nat Commun 12:2862–12. doi:10.1038/s41467-021-22922-6

Bae S-A, Fang MZ, Rustgi V, Zarbl H, Androulakis IP. 2019. At the Interface of Lifestyle, Behavior, and Circadian Rhythms: Metabolic Implications. Front Nutr 6:132. doi:10.3389/fnut.2019.00132

Bargiello TA, Jackson FR, Young MW. 1984. Restoration of circadian behavioural rhythms by gene transfer in Drosophila. Nature 312:752–754.

Bischof J, Maeda RK, Hediger M, Karch F, Basler K. 2007. An optimized transgenesis system for Drosophila using germ-line-specific phiC31 integrases. Proc Natl Acad Sci USA 104:3312–3317. doi:10.1073/pnas.0611511104

Blanchard FJ, Collins B, Cyran SA, Hancock DH, Taylor MV, Blau J. 2010. The transcription factor Mef2 is required for normal circadian behavior in Drosophila. Journal of Neuroscience 30:5855–5865. doi:10.1523/jneurosci.2688-09.2010

Blau J, Young MW. 1999. Cycling vrille expression is required for a functional Drosophila clock. Cell 99:661–671.

Bonnay F, Cohen-Berros E, Hoffmann M, Kim SY, Boulianne GL, Hoffmann JA, Matt N, Reichhart J-M. 2013. big bang gene modulates gut immune tolerance in Drosophila. Proc Natl Acad Sci USA 110:2957–2962. doi:10.1073/pnas.1221910110

Brandes C, Plautz JD, Stanewsky R, Jamison CF, Straume M, Wood KV, Kay SA, Hall JC. 1996. Novel features of drosophila period Transcription revealed by real-time luciferase reporting. Neuron 16:687–692.

Brown AJ, Pendergast JS, Yamazaki S. 2019. Peripheral Circadian Oscillators. Yale J Biol Med 92:327–335. doi:10.1126/sciadv.abg5174

Cripps RM, Black BL, Zhao B, Lien CL, Schulz RA, Olson EN. 1998. The myogenic regulatory gene Mef2 is a direct target for transcriptional activation by Twist during Drosophila myogenesis. Genes & Development 12:422–434. doi:10.1101/gad.12.3.422

Croker B, Crozat K, Berger M, Xia Y, Sovath S, Schaffer L, Eleftherianos I, Imler J-L, Beutler 2007. ATP-sensitive potassium channels mediate survival during infection in mammals and insects. Nat Genet 39:1453–1460. doi:10.1038/ng.2007.25

Erion R, King AN, Wu G, Hogenesch JB, Sehgal A. 2016. Neural clocks and Neuropeptide F/Y regulate circadian gene expression in a peripheral metabolic tissue. Elife 5:e13552. doi:10.7554/elife.13552

Evans CJ, Olson JM, Ngo KT, Kim E, Lee NE, Kuoy E, Patananan AN, Sitz D, Tran P, Do M-T, Yackle K, Cespedes A, Hartenstein V, Call GB, Banerjee U. 2009. G-TRACE: rapid Gal4-based cell lineage analysis in Drosophila. Nature Publishing Group 6:603–605. doi:10.1038/nmeth.1356

Franco DL, Frenkel L, Ceriani MF. 2018. The Underlying Genetics of Drosophila Circadian Behaviors. Physiology (Bethesda) 33:50–62. doi:10.1152/physiol.00020.2017

Gajewski K, Kim Y, Lee YM, Olson EN, Schulz RA. 1997. D-mef2 is a target for Tinman activation during Drosophila heart development. EMBO J 16:515–522. doi:10.1093/emboj/16.3.515

Gill S, Le HD, Melkani GC, Panda S. 2015. Time-restricted feeding attenuates age-related cardiac decline in Drosophila. Science 347:1265–1269. doi:10.1126/science.1256682

Guo F, Cerullo I, Chen X, Rosbash M. 2014. PDF neuron firing phase-shifts key circadian activity neurons in Drosophila. Elife 3:e02780. doi:10.7554/elife.02780

Guo F, Chen X, Rosbash M. 2017. Temporal calcium profiling of specific circadian neurons in freely moving flies. Proc Natl Acad Sci USA 3:201706608–E8787. doi:10.1073/pnas.1706608114

Guo F, Yu J, Jung HJ, Abruzzi KC, Luo W, Griffith LC, Rosbash M. 2016. Circadian neuron feedback controls the Drosophila sleep–activity profile. Nature 536:1–18. doi:10.1038/nature19097

Handler AM, Konopka RJ. 1979. Transplantation of a circadian pacemaker in Drosophila. Nature 279:236–238.

Helfrich-Förster C. 1995. The period clock gene is expressed in central nervous system neurons which also produce a neuropeptide that reveals the projections of circadian pacemaker cells within the brain of Drosophila melanogaster. Proc Natl Acad Sci USA 92:612–616.

Helfrich-Förster C, Täuber M, Park JH, Mühlig-Versen M, Schneuwly S, Hofbauer A. 2000. Ectopic expression of the neuropeptide pigment-dispersing factor alters behavioral rhythms in Drosophila melanogaster. Journal of Neuroscience 20:3339–3353.

Hood S, Amir S. 2017a. The aging clock: circadian rhythms and later life. J Clin Invest 127:437–446. doi:10.1172/jci90328

Hood S, Amir S. 2017b. Neurodegeneration and the Circadian Clock. Front Aging Neurosci 9:170. doi:10.3389/fnagi.2017.00170

Hrdlicka L, Gibson M, Kiger A, Micchelli C, Schober M, Schöck F, Perrimon N. 2002. Analysis of twenty-four Gal4 lines in Drosophila melanogaster. Genesis 34:51–57. doi:10.1002/gene.10125

Hyun S, Lee Y, Hong S-T, Bang S, Paik D, Kang J, Shin J, Lee J, Jeon K, Hwang S, Bae E, Kim J. 2005. Drosophila GPCR Han is a receptor for the circadian clock neuropeptide PDF. Neuron 48:267–278. doi:10.1016/j.neuron.2005.08.025

Ito C, Tomioka K. 2016. Heterogeneity of the Peripheral Circadian Systems in Drosophila melanogaster: A Review. Front Physiol 7:8. doi:10.3389/fphys.2016.00008

Kaneko H, Head LM, Ling J, Tang X, Liu Y, Hardin PE, Emery P, Hamada FN. 2012. Circadian rhythm of temperature preference and its neural control in Drosophila. Curr Biol 22:1851–1857. doi:10.1016/j.cub.2012.08.006

Lee K-Z, Lestradet M, Socha C, Schirmeier S, Schmitz A, Spenlé C, Lefebvre O, Keime C, Yamba WM, Aoun RB, Liegeois S, Schwab Y, Simon-Assmann P, Dalle F, Ferrandon D. 2016. Enterocyte Purge and Rapid Recovery Is a Resilience Reaction of the Gut Epithelium to Pore-Forming Toxin Attack. Cell Host Microbe 20:716–730. doi:10.1016/j.chom.2016.10.010

Leise TL, Harrington ME. 2011. Wavelet-based time series analysis of circadian rhythms. J Biol Rhythms 26:454–463. doi:10.1177/0748730411416330

Leng Y, Musiek ES, Hu K, Cappuccio FP, Yaffe K. 2019. Association between circadian rhythms and neurodegenerative diseases. Lancet Neurol 18:307–318. doi:10.1016/s1474-4422(18)30461-7

Lin Y, Stormo GD, Taghert PH. 2004. The neuropeptide pigment-dispersing factor coordinates pacemaker interactions in the Drosophila circadian system. Journal of Neuroscience 24:7951–7957. doi:10.1523/jneurosci.2370-04.2004

Logan RW, McClung CA. 2019. Rhythms of life: circadian disruption and brain disorders across the lifespan. Nat Rev Neurosci 20:49–65. doi:10.1038/s41583-018-0088-y

Mertens I, Vandingenen A, Johnson EC, Shafer OT, Li W, Trigg JS, Loof AD, Schoofs L, Taghert PH. 2005. PDF Receptor Signaling in Drosophila Contributes to Both Circadian and Geotactic Behaviors. Neuron 48:213–219. doi:10.1016/j.neuron.2005.09.009

Mezan S, Feuz JD, Deplancke B, Kadener S. 2016. PDF Signaling Is an Integral Part of the Drosophila Circadian Molecular Oscillator. CellReports 17:708–719. doi:10.1016/j.celrep.2016.09.048

Murad A, Emery-Le M, Emery P. 2007. A subset of dorsal neurons modulates circadian behavior and light responses in Drosophila. Neuron 53:689–701. doi:10.1016/j.neuron.2007.01.034

Muraro NI, Ceriani MF. 2015. Acetylcholine from Visual Circuits Modulates the Activity of Arousal Neurons in Drosophila. Journal of Neuroscience 35:16315–16327. doi:10.1523/jneurosci.1571-15.2015

Myers MP, Wager-Smith K, Rothenfluh-Hilfiker A, Young MW. 1996. Light-induced degradation of TIMELESS and entrainment of the Drosophila circadian clock. Science 271:1736–1740.

Nern A, Pfeiffer BD, Svoboda K, Rubin GM. 2011. Multiple new site-specific recombinases for use in manipulating animal genomes. Proc Natl Acad Sci USA 108:14198–14203. doi:10.1073/pnas.1111704108

Paredes JC, Welchman DP, Poidevin M, Lemaitre B. 2011. Negative regulation by amidase PGRPs shapes the Drosophila antibacterial response and protects the fly from innocuous infection. Immunity 35:770–779. doi:10.1016/j.immuni.2011.09.018

Park JH, Helfrich-Förster C, Lee G, Liu L, Rosbash M, Hall JC. 2000. Differential regulation of circadian pacemaker output by separate clock genes in Drosophila. Proc Natl Acad Sci USA 97:3608–3613. doi:10.1073/pnas.070036197

Patke A, Young MW, Axelrod S. 2020. Molecular mechanisms and physiological importance of circadian rhythms. Nat Rev Mol Cell Biol 21:67–84. doi:10.1038/s41580-019-0179-2

Picot M, Cusumano P, Klarsfeld A, Ueda R, Rouyer F. 2007. Light activates output from evening neurons and inhibits output from morning neurons in the Drosophila circadian clock. PLoS Biol 5:e315. doi:10.1371/journal.pbio.0050315

Pilorz V, Helfrich-Förster C, Oster H. 2018. The role of the circadian clock system in physiology. Pflugers Arch 470:227–239. doi:10.1007/s00424-017-2103-y

Rana S, Prabhu SD, Young ME. 2020. Chronobiological Influence Over Cardiovascular Function: The Good, the Bad, and the Ugly. Circ Res 126:258–279. doi:10.1161/circresaha.119.313349

Renn SC, Park JH, Rosbash M, Hall JC, Taghert PH. 1999. A pdf neuropeptide gene mutation and ablation of PDF neurons each cause severe abnormalities of behavioral circadian rhythms in Drosophila. Cell 99:791–802.

Roberts L, Leise TL, Noguchi T, Galschiodt AM, Houl JH, Welsh DK, Holmes TC. 2015. Light evokes rapid circadian network oscillator desynchrony followed by gradual phase retuning of synchrony. Curr Biol 25:858–867. doi:10.1016/j.cub.2015.01.056

Rothenfluh A, Abodeely M, Price JL, Young MW. 2000a. Isolation and analysis of six timeless alleles that cause short- or long-period circadian rhythms in Drosophila. Genetics 156:665–675.

Rothenfluh A, Young MW, Saez L. 2000b. A TIMELESS-independent function for PERIOD proteins in the Drosophila clock. Neuron 26:505–514. doi:10.1016/s0896-6273(00)81182-4

Saitoe M, Schwarz TL, Umbach JA, Gundersen CB, Kidokoro Y. 2001. Absence of junctional glutamate receptor clusters in Drosophila mutants lacking spontaneous transmitter release. Science 293:514–517. doi:10.1126/science.1061270

Schindelin J, Arganda-Carreras I, Frise E, Kaynig V, Longair M, Pietzsch T, Preibisch S, Rueden C, Saalfeld S, Schmid B, Tinevez J-Y, White DJ, Hartenstein V, Eliceiri K, Tomancak P, Cardona A. 2012. Fiji: an open-source platform for biological-image analysis. Nature Publishing Group 9:676–682. doi:10.1038/nmeth.2019

Schmid B, Helfrich-Förster C, Yoshii T. 2011. A new ImageJ plug-in “ActogramJ” for chronobiological analyses. J Biol Rhythms 26:464–467. doi:10.1177/0748730411414264

Schuster CM, Davis GW, Fetter RD, Goodman CS. 1996. Genetic dissection of structural and functional components of synaptic plasticity. I. Fasciclin II controls synaptic stabilization and growth. Neuron 17:641–654. doi:10.1016/s0896-6273(00)80197-x

Selcho M, Millán C, Palacios-Muñoz A, Ruf F, Ubillo L, Chen J, Bergmann G, Ito C, Silva V, Wegener C, Ewer J. 2017. Central and peripheral clocks are coupled by a neuropeptide pathway in Drosophila. Nat Commun 8:15563. doi:10.1038/ncomms15563

Shafer OT, Kim DJ, Dunbar-Yaffe R, Nikolaev VO, Lohse MJ, Taghert PH. 2008. Widespread receptivity to neuropeptide PDF throughout the neuronal circadian clock network of Drosophila revealed by real-time cyclic AMP imaging. Neuron 58:223–237. doi:10.1016/j.neuron.2008.02.018

Shafer OT, Rosbash M, Truman JW. 2002. Sequential nuclear accumulation of the clock proteins period and timeless in the pacemaker neurons of Drosophila melanogaster. Journal of Neuroscience 22:5946–5954.

Shimizu I, Yoshida Y, Minamino T. 2016. A role for circadian clock in metabolic disease. Hypertens Res 39:483–491. doi:10.1038/hr.2016.12

Sink H, Rehm EJ, Richstone L, Bulls YM, Goodman CS. 2001. sidestep encodes a target-derived attractant essential for motor axon guidance in Drosophila. Cell 105:57–67. doi:10.1016/s0092-8674(01)00296-3

So WV, Rosbash M. 1997. Post-transcriptional regulation contributes to Drosophila clock gene mRNA cycling. EMBO J 16:7146–7155. doi:10.1093/emboj/16.23.7146

Stanewsky R, Jamison CF, Plautz JD, Kay SA, Hall JC. 1997. Multiple circadian-regulated elements contribute to cycling period gene expression in Drosophila. EMBO J 16:5006– 5018. doi:10.1093/emboj/16.16.5006

Strand M, Micchelli CA. 2011. Quiescent gastric stem cells maintain the adult Drosophila stomach. Proc Natl Acad Sci USA 108:17696–17701. doi:10.1073/pnas.1109794108

Sulli G, Lam MTY, Panda S. 2019. Interplay between Circadian Clock and Cancer: New Frontiers for Cancer Treatment. Trends Cancer 5:475–494. doi:10.1016/j.trecan.2019.07.002

Takeuchi T, Suzuki M, Fujikake N, Popiel HA, Kikuchi H, Futaki S, Wada K, Nagai Y. 2015. Intercellular chaperone transmission via exosomes contributes to maintenance of protein homeostasis at the organismal level. Proc Natl Acad Sci USA 112:E2497–506. doi:10.1073/pnas.1412651112

Tanenhaus AK, Zhang J, Yin JCP. 2012. In Vivo Circadian Oscillation of dCREB2 and NF-κB Activity in the Drosophila Nervous System. Plos One 7:e45130. doi:10.1371/journal.pone.0045130

Thosar SS, Butler MP, Shea SA. 2018. Role of the circadian system in cardiovascular disease. J Clin Invest 128:2157–2167. doi:10.1172/jci80590

Top D, Harms E, Syed S, Adams EL, Saez L. 2016. GSK-3 and CK2 Kinases Converge on Timeless to Regulate the Master Clock. CellReports 16:357–367. doi:10.1016/j.celrep.2016.06.005

Top D, O’Neil JL, Merz GE, Dusad K, Crane BR, Young MW. 2018. CK1/Doubletime activity delays transcription activation in the circadian clock. Elife 7:e32679. doi:10.7554/elife.32679

Tsuchiya Y, Umemura Y, Yagita K. 2020. Circadian clock and cancer: From a viewpoint of cellular differentiation. Int J Urol 27:518–524. doi:10.1111/iju.14231

Veleri S, Wülbeck C. 2004. Unique self-sustaining circadian oscillators within the brain of Drosophila melanogaster. Chronobiol Int 21:329–342. doi:10.1081/cbi-120038597

Versteven M, Ernst K-M, Stanewsky R. 2020. A Robust and Self-Sustained Peripheral Circadian Oscillator Reveals Differences in Temperature Compensation Properties with Central Brain Clocks. iScience 23:101388. doi:10.1016/j.isci.2020.101388

Yang Z, Sehgal A. 2001. Role of molecular oscillations in generating behavioral rhythms in Drosophila. Neuron 29:453–467.

Yoshii T, Wülbeck C, Sehadova H, Veleri S, Bichler D, Stanewsky R, Helfrich-Förster C. 2009. The neuropeptide pigment-dispersing factor adjusts period and phase of Drosophila’s clock. Journal of Neuroscience 29:2597–2610. doi:10.1523/jneurosci.5439-08.2009

Zaidman-Rémy A, Hervé M, Poidevin M, Pili-Floury S, Kim M-S, Blanot D, Oh B-H, Ueda R, Mengin-Lecreulx D, Lemaitre B. 2006. The Drosophila amidase PGRP-LB modulates the immune response to bacterial infection. Immunity 24:463–473. doi:10.1016/j.immuni.2006.02.012

Zhang L, Chung BY, Lear BC, Kilman VL, Liu Y, Mahesh G, Meissner R-A, Hardin PE, Allada R. 2010. DN1(p) circadian neurons coordinate acute light and PDF inputs to produce robust daily behavior in Drosophila. Curr Biol 20:591–599. doi:10.1016/j.cub.2010.02.056

Zhang Y, Liu Y, Bilodeau-Wentworth D, Hardin PE, Emery P. 2010. Light and temperature control the contribution of specific DN1 neurons to Drosophila circadian behavior. Curr Biol 20:600–605. doi:10.1016/j.cub.2010.02.044

Zhang Z, Zeng P, Gao W, Zhou Q, Feng T, Tian X. 2021. Circadian clock: a regulator of the immunity in cancer. Cell Communication and Signaling 19:37. doi:10.1186/s12964-021-00721-2

