## Supplementary figures and images for "Real time, *in vivo* measurement of neuronal and peripheral clocks in *Drosophila melanogaster*"

### Supplemental Figure 1

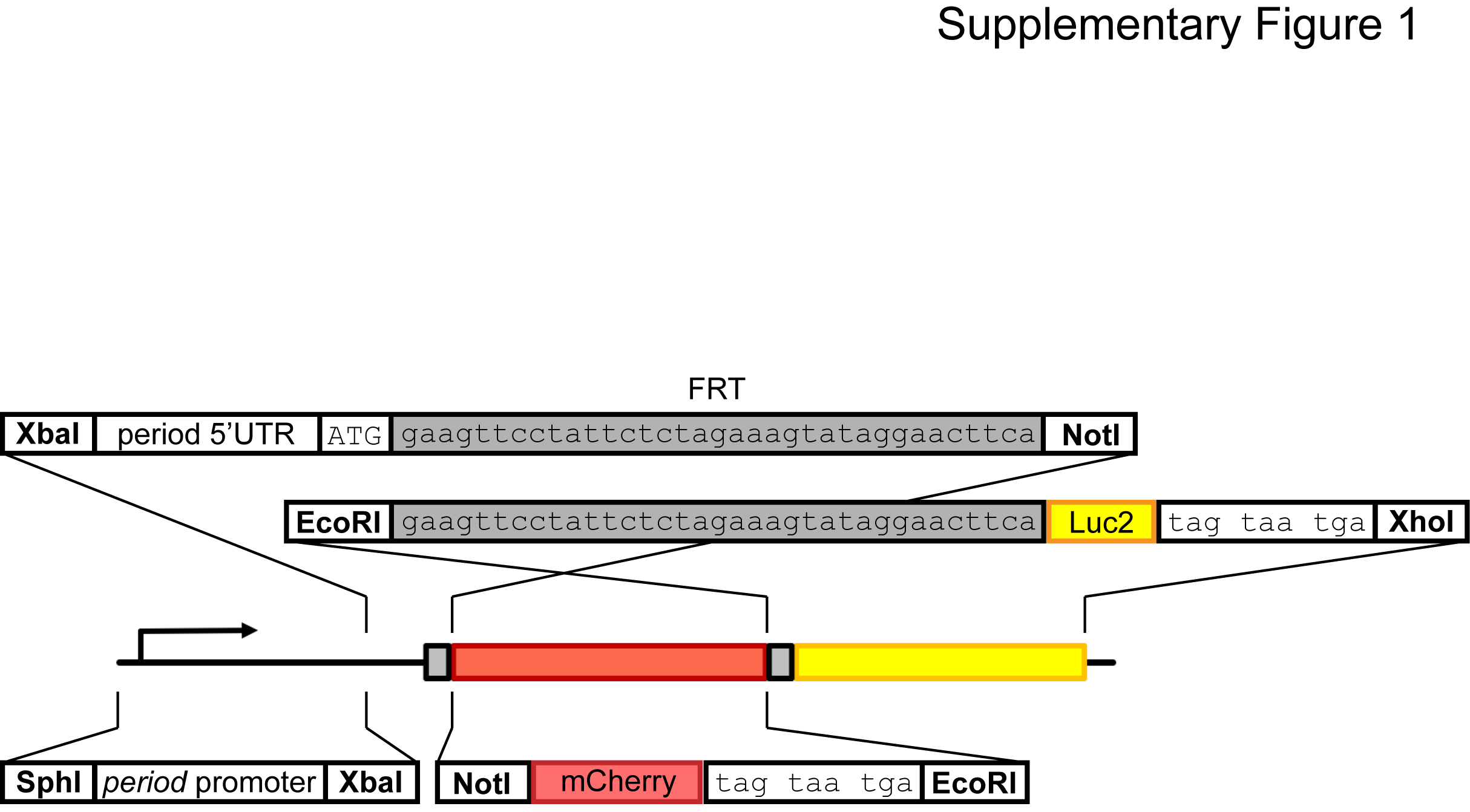

### Supplemental Figure 2

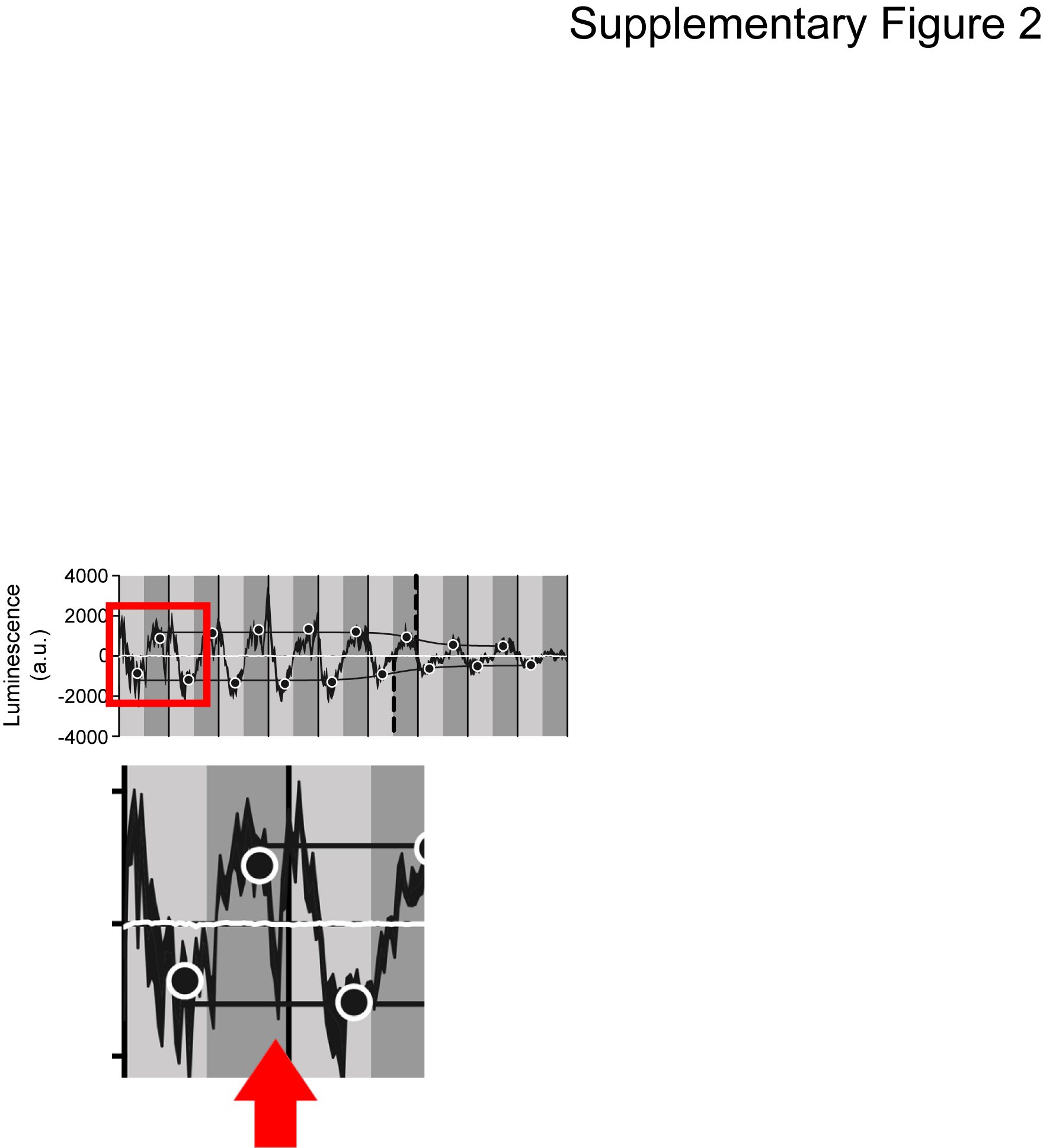
