## Supplemental Table 1 for "Real time, *in vivo* measurement of neuronal and peripheral clocks in *Drosophila melanogaster*"

### SUPPLEMENTARY FIGURE LEGENDS

**Supplementary Table 1.** Genotypes of flies used in the figures. Stocks are maintained with one parental line expressing both the Gal4 driver and LABL reporter, and the other parental line expressing UAS-*flp*.

| Figure | Label | Genotype |
|---|---|---|
| Figure 2 | tim-UAS-Gal4 | w ; tim-UAS-Gal4 / + ; LABL / UAS-flp2 |
| Figure 3A | tim-UAS-Gal4 | w ; tim-UAS-Gal4 / + ; LABL / UAS-flp2 |
| Figure 3A | plo | w ; plo ; + |
| Figure 3A | PER-BG::Luc | w ; per-BG-luc ; + |
| Figure 3B | $tim^{01}$ | w ; tim-UAS-Gal4, $tim^{01}$ / $tim^{01}$ ; LABL / UAS-flp2 |
| Figure 3C | $han^{5304}$ | w,$han^{5304}$ ; tim-UAS-Gal4 / + ; LABL / UAS-flp2 |
| Figure 4B-4C, 5A-5D | Pdf-Gal4 | w ; Pdf-Gal4 / + ; LABL / UAS-flp2 |
| Figure 4B-4C, 5A-5D | DvPdf-Gal4 | w ; DvPdf-Gal4 / + ; LABL / UAS-flp2 |
| Figure 4B-4C, 5A-5D | R18H11-Gal4 | w ; LABL / + ; R18H11-Gal4 / UAS-flp2 |
| Figure 4B-4C, 5A-5D | tim-UAS-Gal4 | w ; tim-UAS-Gal4 / + ; LABL / UAS-flp2 |
| Figure 4B, 5A-5D | Clk4.1-Gal4 | w ; LABL / + ; Clk4.1-Gal4 / UAS-flp2 |
| Figure 4B, 5A-5D | Mai179-Gal4 | w ; Mai179-Gal4 / + ; LABL / UAS-flp2 |
| Figure 4B, 5A-5D | Clk9M-Gal4 | w ; Clk9M-Gal4 / + ; LABL / UAS-flp2 |
| Figure 4D | Pdf-Gal4 | w ; Pdf-Gal4 / + ; + / G-TRACE |
| Figure 4D | DvPdf-Gal4 | w ; DvPdf-Gal4 / + ; + / G-TRACE |
| Figure 4D | R18H11-Gal4 | w ; + / + ; R18H11 / G-TRACE |
| Figure 5A-5D | tim-UAS-Gal4 \| $han^{5304}$ | $han^{5304}$ ; tim-UAS-Gal4 / + ; LABL / UAS-flp2 |
| Figure 5A-5D | Pdf \| $han^{5304}$ | $han^{5304}$ ; Pdf-Gal4 / + ; LABL / UAS-flp2 |
| Figure 5A-5D | DvPdf \| $han^{5304}$ | $han^{5304}$ ; DvPdf-Gal4 / + ; LABL / UAS-flp2 |
| Figure 5A-5D | R18H11 \| $han^{5304}$ | $han^{5304}$ ; LABL / + ; R18H11-Gal4 / UAS-flp2 |
| Figure 5A-5D | Clk4.1 \| $han^{5304}$ | $han^{5304}$ ; LABL / + ; Clk4.1-Gal4 / UAS-flp2 |
| Figure 5A-5D | Mai179 \| $han^{5304}$ | $han^{5304}$ ; Mai179-Gal4 / + ; LABL / UAS-flp2 |
| Figure 5A-5D | Clk9M \| $han^{5304}$ | $han^{5304}$ ; Clk9M-Gal4 / + ; LABL / UAS-flp2 |
| Figure 6B-6E | elav-Gal4 | elav-Gal4 ; + / + ; LABL / UAS-flp2 |
| Figure 6B-6E | mef2-Gal4 | w ; LABL / + ; mef2-Gal4 / UAS-flp2 |
| Figure 6B-6E | esg-Gal4 | w ; esg-Gal4 / + ; LABL / UAS-flp2 |
| Figure 6B-6E | NP3084-Gal4 | w ; LABL / + ; NP3084-Gal4 / UAS-flp2 |
| Figure 6B-6E | C564-Gal4 | w ; C564-Gal4 / + ; LABL / UAS-flp2 |
| Figure 6B-6E | LSP2-Gal4 | w ; LABL / + ; LSP2-Gal4 / UAS-flp2 |
| Figure 6B-6E | mef2-Gal4 \| $han^{5304}$ | $han^{5304}$ ; LABL / + ; mef2-Gal4 / UAS-flp2 |
| Figure 6B-6E | esg-Gal4 \| $han^{5304}$ | $han^{5304}$ ; esg-Gal4 / + ; LABL / UAS-flp2 |
| Figure 6B-6E | NP3084-Gal4 \| $han^{5304}$ | $han^{5304}$ ; LABL / + ; NP3084-Gal4 / UAS-flp2 |
| Figure 6B-6E | C564-Gal4 \| $han^{5304}$ | $han^{5304}$ ; C564-Gal4 / + ; LABL / UAS-flp2 |
| Figure 6B-6E | LSP2-Gal4 \| $han^{5304}$ | $han^{5304}$ ; LABL / + ; LSP2-Gal4 / UAS-flp2 |
